# SARS-CoV-2 Exploits Sexually Dimorphic and Adaptive IFN and TNFa Signaling to Gain Entry into Alveolar Epithelium

**DOI:** 10.1101/2021.07.23.453505

**Authors:** Yan Wang, Sreeharsha Gurrapu, Hong Chen, Sara Laudato, Emily Caggiano, Yan Jiang, Hsiang-Hsi Ling, Filippo G. Giancotti

**Author notes:** These authors contributed equally to this work (Yan Wang, Sreeharsha Gurrapu, Hong Chen, Sara Laudato).

## Abstract

Infection of the alveolar epithelium constitutes a bottleneck in the progression of COVID-19 to SARS presumably due to the paucity of viral entry receptors in alveolar epithelial type 1 and 2 cells. We have found that the male alveolar epithelial cells express twice as many ACE2 and TMPRSS2 entry receptors as the female ones. Intriguingly, IFN and TNF-α signaling are preferentially active in male alveolar cells and induce binding of the cognate transcription factors to the promoters and lung-active enhancers of *ACE2* and *TMPRSS2*. Cotreatment with IFN-I and III dramatically increases expression of the receptors and viral entry in alveolar epithelial cells. TNFα and IFN-II, typically overproduced during the cytokine storm, similarly collaborate to induce these events. Whereas JAK inhibitors suppress viral entry induced by IFN-I/III, simultaneous inhibition of IKK/NF-*κ*B is necessary to block viral entry induced by TNFα and IFN-II. In addition to explaining the increased incidence of SARS in males, these findings indicate that SARS-Cov-2 hijacks epithelial immune signaling to promote infection of the alveolar epithelium and suggest that JAK inhibitors, singly and in combination with NF-KB inhibitors, may exhibit efficacy in preventing or treating COVID-19 SARS.

## Introduction

The Coronavirus Disease-19 (COVID-19) pandemic has illustrated the power of novel airborne viruses to spread rapidly amongst immunologically naïve human populations and cause severe lung disease and death in individuals with predisposing conditions (1). Although public health measures, such as lockdowns, have been effective, they have imposed a heavy economic and social price, which has limited their full implementation (2). Extraordinary efforts have led to the recent development of safe and effective vaccines, and large-scale vaccination efforts are underway in several major countries (3). However, it remains uncertain whether these efforts will be so successful to eradicate the disease (4, 5). In order to develop better therapeutics for COVID-19 and novel coronaviruses that may emerge in the future, it is important to understand the biology of viral infection, viral pathogenicity, and immune response to the disease.

The etiological agent of COVID-19, Severe Acute Respiratory Syndrome Coronavirus-2 (SARS-CoV-2), is an enveloped RNA virus decorated by spike (S) protein trimers, which mediate entry into host cells (6). Although SARS-Cov-2 usually produces only a mild upper respiratory tract infection, it spreads to the distal lung in a minor but significant fraction of individuals, causing bilateral pneumonia characterized by severe hypoxia (7). Simultaneous or consequent systemic dissemination underlies extrapulmonary manifestations of the disease, including multiorgan injury, endothelial cell damage and ensuing micro thromboembolism, and severe dysregulation of immune responses (8). These observations suggest that viral entry into and damage to the alveolar epithelium occurs infrequently but is a major determinant of progression to severe disease.

The entry of SARS-Cov-2 into target cells requires proteolytic activation of the S protein mediated by the cell surface serine protease TMPRSS2 and subsequent binding of cleaved, fusogenic S to the entry receptor Angiotensin-Converting Enzyme 2 (ACE2) (9). In contrast to what previously assumed, viral exit is not mediated by the secretory pathway, but by lysosomal trafficking (10). This non-lytic release mechanism is associated with lysosome deacidification and inactivation of lysosomal enzymes, making it unlikely to generate a virus with activated S protein. These considerations suggest that SARS-Cov-2 entry into cells is critically dependent on the co-expression of ACE2 and TMPRSS2 in target cells.

Reverse genetics experiments have revealed a proximal to distal gradient in sensitivity to infection of the airway epithelium, which correlates with decreasing levels of expression of ACE2, but not TMPRSS2 (11). Single-cell RNA sequencing (scRNA-seq) analysis has indicated that goblet and ciliated cells from the nasal epithelium express elevated levels of ACE2 and TMPRSS2, suggesting that they constitute the initial entry points for infection (12). Although ACE2 and TMPRSS2 are expressed in bronchial secretory cells, where ACE2 is upregulated by smoking via inflammatory signaling (13, 14), ACE2 and TMPRSS2 are co-expressed only in a small subset of alveolar epithelial type II cells, suggesting that viral entry may require upregulation of ACE2 or both ACE2 and TMPRSS2 (11, 15).

The mechanisms that regulate the expression of ACE2 and TMPRSS2 in lung alveolar epithelium are incompletely understood. Expression of ACE2 correlates with elevated interferon (IFN) signaling in a subset of type II alveolar epithelial cells (15). In addition, it has been reported that treatment with IFNαinduces upregulation of ACE2 in stem cell-derived alveolo-spheres consisting of type II cells (16). However, recent findings have indicated that IFN signaling only induces expression of a truncated form of ACE2, which lacks the N-terminal S protein-binding site and is generated through cooption of an IFN-responsive endogenous retroelement (17, 18). In parallel, association studies have lent support to the notion that TMPRSS2 is upregulated by androgen receptor (AR) signaling in lung epithelium as it is in the normal prostate and androgen-dependent prostate cancer (19). In fact, prostate tumorigenesis is often initiated by oncogenic fusions bringing an oncogenic ETS factor, such as ERG or ETV1, under the control of the AR-regulated promoter of *TMPRSS2* (20). Consistently, a recent study has shown that dutasteride, an androgen biosynthesis inhibitor, inhibits the expression of *TMPRSS2* in alveolar epithelial cells (21). In addition, preliminary evidence suggests that the incidence of severe COVID-19 disease is reduced in prostate cancer patients treated with second-generation AR inhibitors, such as enzalutamide and abiraterone (22). However, it has not been yet shown if blockade of the AR with enzalutamide or other direct inhibitors reduces TMPRSS2 expression in alveolar epithelial cells.

Early studies have indicated that the age-adjusted incidence and mortality of COVID-19 are strikingly higher in males as compared to females (23, 24). Patients progressing to severe disease exhibit defective B and T cell responses (25, 26) and a delayed but exaggerated activation of innate immune system, culminating in overproduction of multiple cytokines and systemic toxicity (27–29). Intriguingly, higher plasma levels of innate immune cytokines and non-classical monocytes, but poorer T cell responses, are observed in men as compared to women during moderate disease, suggesting that sex differences in immune responses may contribute to the higher incidence of severe disease and death in men (30). Although it is known that women mount stronger immune responses against viruses and vaccines and exhibit superior immune-mediated tissue repair (31, 32), the molecular and cellular mechanisms underlying sex-differences in immunity and their relationship to SARS-CoV-2 pathogenesis are not well understood (33).

In this study, we have examined the role of innate and adaptive IFN and NF-*κ*B signaling in regulating the expression of viral entry receptors and infection of alveolar epithelium in COVID-19. Our findings indicate that sexually dimorphic IFN signaling upregulates the expression of *ACE2* and *TMPRSS2* in the alveolar epithelium of males, potentially explaining the higher incidence of SARS in this gender. In addition, we provide evidence that SARS-Cov-2 hijacks the innate IFN-I and III response and the adaptive IFN-II and NF-*κ*B response to promote its entry into target cells. We propose that rational combinations of a JAK inhibitor and antivirals may prevent progression to SARS, whereas combinations of a JAK inhibitor and a NF-*κ*B inhibitor may exhibit therapeutic efficacy in advanced stage SARS.

## Results

### *ACE2* and *TMPRSS2* are Co-Expressed in a Subset of Type I and II Alveolar Epithelial Cells in Normal Individuals

To examine if the expression of the canonical SARS-CoV-2 entry factors *ACE2* and *TMPRSS2* is sexually dimorphic in normal adult lung, we merged three single-cell RNA-Seq datasets for which gender information was available (14, 34, 35) (Fig. 1A). In total, we analyzed 93,770 single-cell transcriptomes from 9 males and 15 females. Data were integrated by using Harmony, an algorithm that projects cells into a shared embedding in which cells are grouped by cell type rather than dataset-specific conditions (36). Samples were finally subjected to unsupervised graph-based clustering (Fig. 1A).

**Figure 1.**
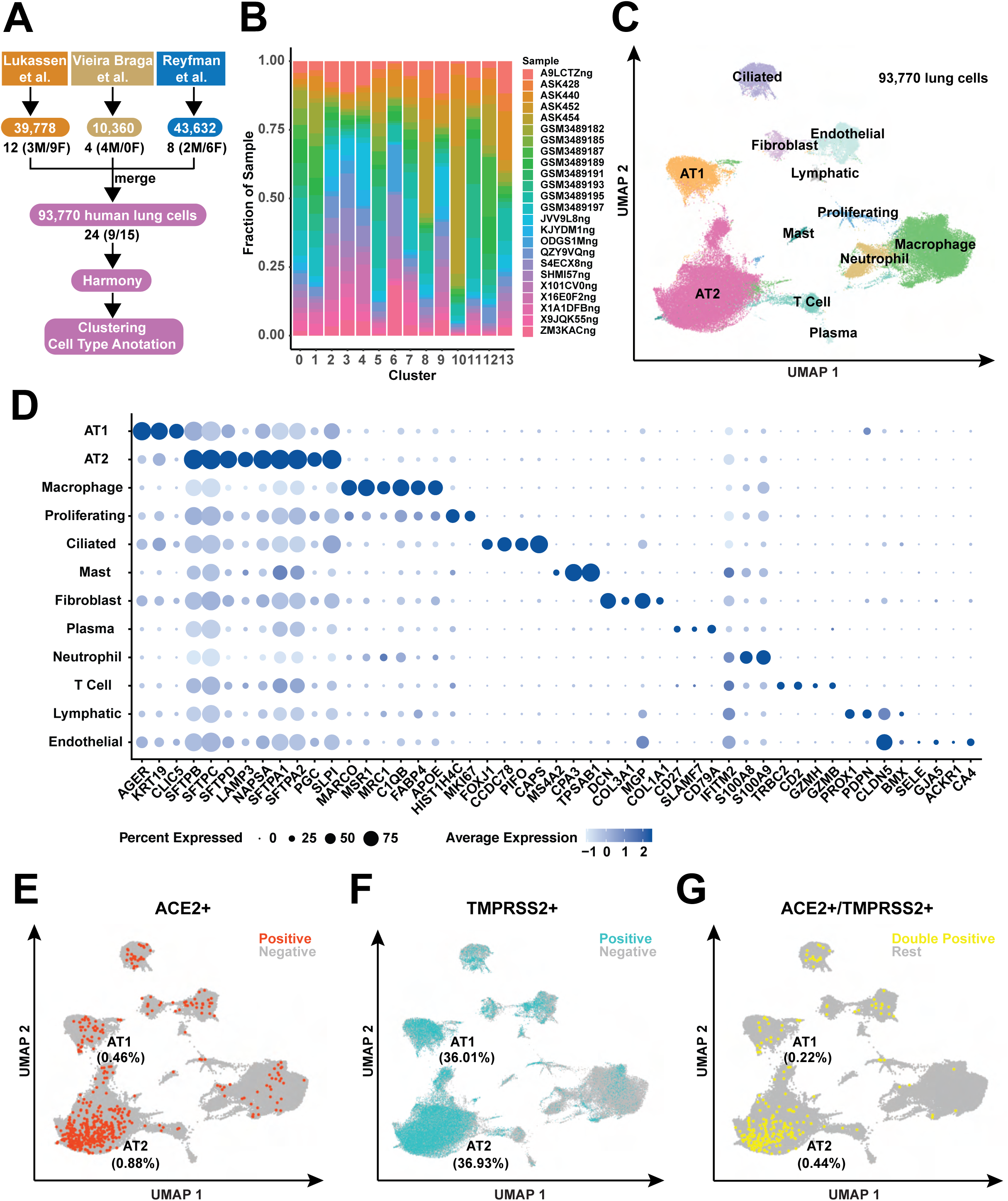
Gene expression of ACE2 and TMPRSS2 occurs largely in AT1 and AT2 epithelial cells. (A) Schematic demonstrating the integration of multiple single cell RNA sequencing datasets from healthy human lung samples (n = 24). (B) Stacked bar plot showing the fraction of each sample represented in each cluster. X-axis is the cluster (#0-13), and y-axis is the fraction of each sample that makes up each cluster. (C) UMAP visualization of 93,770 human lung cells colored by cell type and annotated on the basis of marker genes. (D) Expression of selected canonical cell markers by cell type. The size of the dot correlates to the percentage of cells within a cell type in which that marker was detected. The color shows the average expression level. AT1 = alveolar type I; AT2 = alveolar type II. (E-G) UMAP projection with cells colored by detection of *ACE2* (E), *TMPRSS2* (F) or *ACE2+/TMPRSS2+* (G). The percentages of AT1 and of AT2 cells expressing *ACE2*, *TMPRSS2* or *ACE2/TMPRSS2*, respectively, are also given.

Uniform Manifold Approximation and Projection for Dimension Reduction (UMAP) revealed that the lung transcriptomes from 24 individuals aggregate in 14 clusters (0–13) (fig. S1A). Attesting to successful batch correction, there was limited variability in the cluster distribution of single-cell transcriptomes from the 24 individuals (Fig. 1B). Moreover, the entropy of mixing the sample batches approximated that of negative controls (fig. S1B). UMAP visualization and phylogenetic analysis of identity classes using BuildClusterTree indicated that clusters 0 and 3 and clusters 1 and 7 are closely related (fig. S1C and S1D). Analysis of the expression of canonical markers of each cell type in the normal lung confirmed this observation (fig. S1E). Merging the data in these 2 pairs of clusters led to the definition of 12 clusters. Based on the expression of marker genes, these clusters were annotated as Alveolar Type I (AT1) and II (AT2) cells, ciliated cells, fibroblasts, endothelial cells from blood vessels or lymphatic vessels, macrophages, neutrophils, mast cells, plasma cells, and proliferating cells (Fig. 1C and 1D).

TMPRSS2-mediated cleavage is required for priming the spike S protein of SARS-Cov-2 so that it can bind to ACE2 and mediate membrane fusion (9). Analysis of the scRNA-Seq dataset indicated that *ACE2* is expressed only on a small proportion of AT1 and AT2 cells in the normal lung (∼0.5% and ∼0.9%, respectively) (Fig. 1E), as anticipated from reverse genetics studies (11). In contrast, *TMPRSS2* is expressed in a similarly sizeable fraction of AT1 and AT2 cells (∼36% and ∼37%, respectively) (Fig. 1F). Intriguingly, cells co-expressing *ACE2* and *TMPRSS2* constituted only a minority of the AT1, AT2, ciliated, and endothelial subsets (Fig. 1G). In fact, only 0.22% of AT1 and 0.44% of AT2 cells co-expressed the SARS-CoV-2 viral entry receptors. These results suggest that progression to SARS is constrained by the limiting number of alveolar epithelial cells co-expressing viral entry receptors and point to the existence of signaling mechanisms that elevate their expression.

### The Expression of *ACE2* and *TMPRSS2* is Significantly Higher in Male AT1 and AT2 Cells as Compared to their Female Counterparts

Considering the larger incidence of SARS in men as compared to women, we examined the distribution and level of expression of *ACE2* and *TMPRSS2* in the epithelial and non-epithelial compartments of the lung in the two genders. A preliminary analysis indicated that the normal male and female lungs contain similarly sized subpopulations of epithelial and non-epithelial subsets of cells (fig. S2A). Direct comparison of transcriptional data revealed that the percentage and level of expression of *ACE2* in AT2 and AT1 cells and *TMPRSS2* in AT1 cells are larger in males as compared to females (Fig. 2A and 2B). In contrast, male ciliated bronchial cells expressed lower levels of both *ACE2* and *TMPRSS2* as compared to their female counterpart, suggesting that the enrichment of viral entry receptors in the lung epithelium of males is specific to the alveolar epithelial cells (Fig. 2A and 2B). In consonance with this observation, further analysis indicated that the average level of expression of *ACE2* in individual AT1 and AT2 cells and *TMPRSS2* in AT1 cells is significantly higher in males as compared to females (Fig. 2C and 2D). Although we observed a trend towards elevated expression of *TMPRSS2* also in male AT2 cells, the difference was not statistically significant (Fig. 2D). Finally, drawing from these differences in the percentage and level of expression of *ACE2* and *TMPRSS2*, we found that the alveolar epithelium of males contains approximately twice as many ACE2+ TMPRSS2+ double-positive AT1 and AT2 cells as compared to its female counterpart (Fig. 2E). In agreement with prior observations, smokers possessed a larger number of ACE2+ TMPRSS2+ double-positive AT2 cells as compared to non-smokers (fig. S2B) (37). In addition, individuals aged 65 or more also exhibited more double-positive AT1 and AT2 cells, but the results did not reach statistical significance. These observations suggest that the alveolar epithelium of normal adult males contains a significantly larger number of cells potentially sensitive to SARS-Cov-2 infection as compared to its female counterpart.

**Figure 2.**
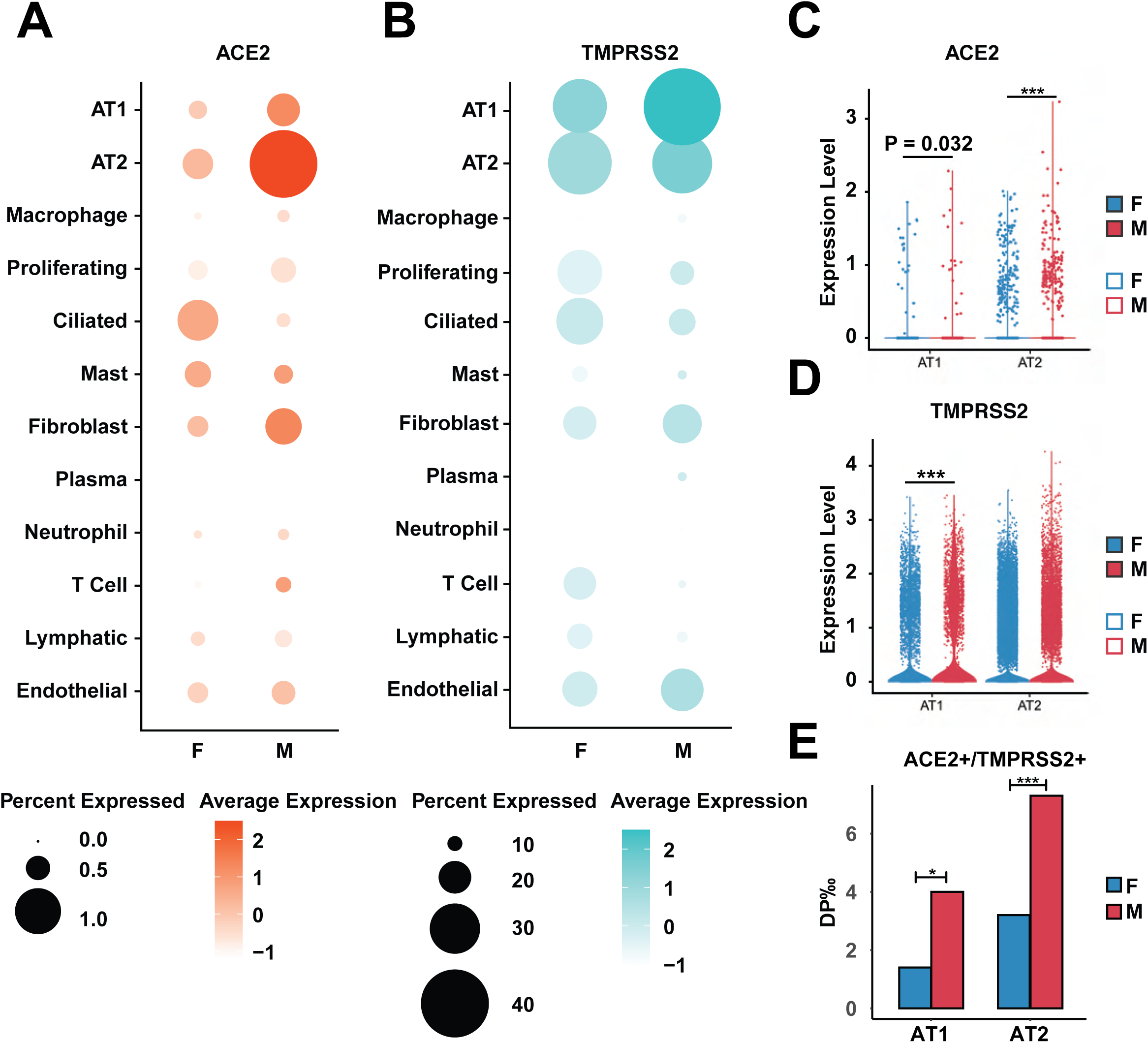
The expression level of ACE2 and TMPRSS2 is sex-related. (A and B) Expression of *ACE2* (A) and *TMPRSS2* (B) in males and females, respectively, by cell type. The size of the dot correlates to the percentage of cells within a cell type in which ACE2/TMPRSS2 was detected. The color encodes the average expression level. (C and D) Normalized expression of *ACE2* (C) and *TMPRSS2* (D) in males (red) and females (blue) in AT1 and AT2 cells. Asterisks indicate significant adjusted p-value of comparison between males and females. *** p.adj < 0.001. (E) The ratio of *ACE2+/TMPRSS2+* double positive cells in AT1 (left) and AT2 (right) cells in males (red) and females (blue). * p < 0.05; *** p < 0.001.

To examine the cell-type specificity of the sexually dimorphic expression of viral entry receptors, we analyzed available single-cell datasets from organs that can be infected by SARS-CoV2 (27, 38–41). The results indicated that the expression of *ACE2* and *TMPRSS2* in epithelial and non-epithelial subsets of the colon, esophagus, and stomach does not vary to a significant degree between sexes (data not shown). We also did not detect gender-based differences in the expression of *ACE2* and *TMPRSS2* in circulating leukocyte subsets (data not shown) (42). Finally, analysis of a single cell dataset from the lungs of patients with COVID-19 (27) indicated that the expression of viral entry receptors is restricted to epithelial cells, macrophages, and T cells (fig. S2C). As previously reported (29, 43), the epithelial compartment of the lungs from COVID-19 patients exhibited elevated expression of ACE2 but not TMPRSS2 as compared to its normal counterpart (fig. S2D and S2E). These observations indicate that the male-predominant expression of *ACE2* and *TMPRSS2* is restricted to the alveolar epithelium.

Recently, it has been proposed that SARS-CoV-2 can enter into certain cell types by combining with a soluble form of ACE2 (sACE2) in the extracellular space. The virus-sACE2 complex would then undergo receptor-mediated endocytosis by binding to vasopressin and thereby to its receptor AVPR1B or by binding directly to the angiotensin II receptor type 1 (AT1, *AGTR1*) (44). Interestingly, scRNA-seq indicated that neither AT1 nor AT2 cells express detectable levels of *AVPRB1* or *AGTR1* (fig. S2F and S2G). Consistently, an analysis of datasets from multiple organs confirmed that these receptors are expressed in the kidney and heart, but not in the lung (NCBI). Finally, qPCR analysis showed that primary human lung alveolar epithelial cells (AEpiC) express low levels of *AVPRB1* or *AGTR1*, in fact, several-fold lower as compared to those of *TMPRSS2* (fig. S2H). These results confirm that the major viral entry receptors on alveolar epithelial cells are *ACE2* and *TMPRSS2* and indicate that their elevated expression in males could engender a higher sensitivity to SARS-CoV2 infection.

### Sexually Dimorphic Interferon Signaling Potentially Controls the Expression of *TMPRSS2* and *ACE2* in Alveolar Epithelial Type I and II Cells

It has been proposed that the AR promotes the expression of *TMPRSS2* in the male lung epithelium, whereas the ER attenuates the expression of *ACE2* in its female counterpart (19, 45, 46). To examine the regulatory regions governing the expression of *TMPRSS2* and *ACE2* in tissues that may be infected by SARS-Cov-2, we examined the Encyclopedia of DNA Elements (ENCODE) compendium. Analysis of the binding profiles for H3K27ac and H3K4me3 identified one upstream enhancer (1) and two distal enhancers (2 and 3) associated with *TMPRSS2* (fig. S3A). Intriguingly, we found that enhancer 1, which corresponds to the classical AR-regulated enhancer active in the prostate gland, is also active in the transverse colon but not in the lung. In contrast, the newly identified enhancers 2 and 3 are active in the lung and, individually, in the thoracic aorta and coronary artery or the liver, respectively (fig. S3A). A similar analysis of *ACE2* revealed a single intronic enhancer active in the lung and transverse colon (fig. S3B). Analysis of the promoter and enhancers of *TMPRSS2* and *ACE2* identified consensus sequences for binding to several transcription factors (TFs), which were ranked based on the best fit for binding. This analysis revealed optimal consensus binding sites for STAT1 (GAS motifs), STAT1/2 (ISRE motifs), various Interferon Response Factors (IRFs), and NF-*κ*B in the regulatory regions of both *TMPRSS2* and *ACE2* (Fig. 3A and 3B), suggesting that IFN and NF-*κ*B signaling regulates the expression of viral entry receptors.

**Figure 3.**
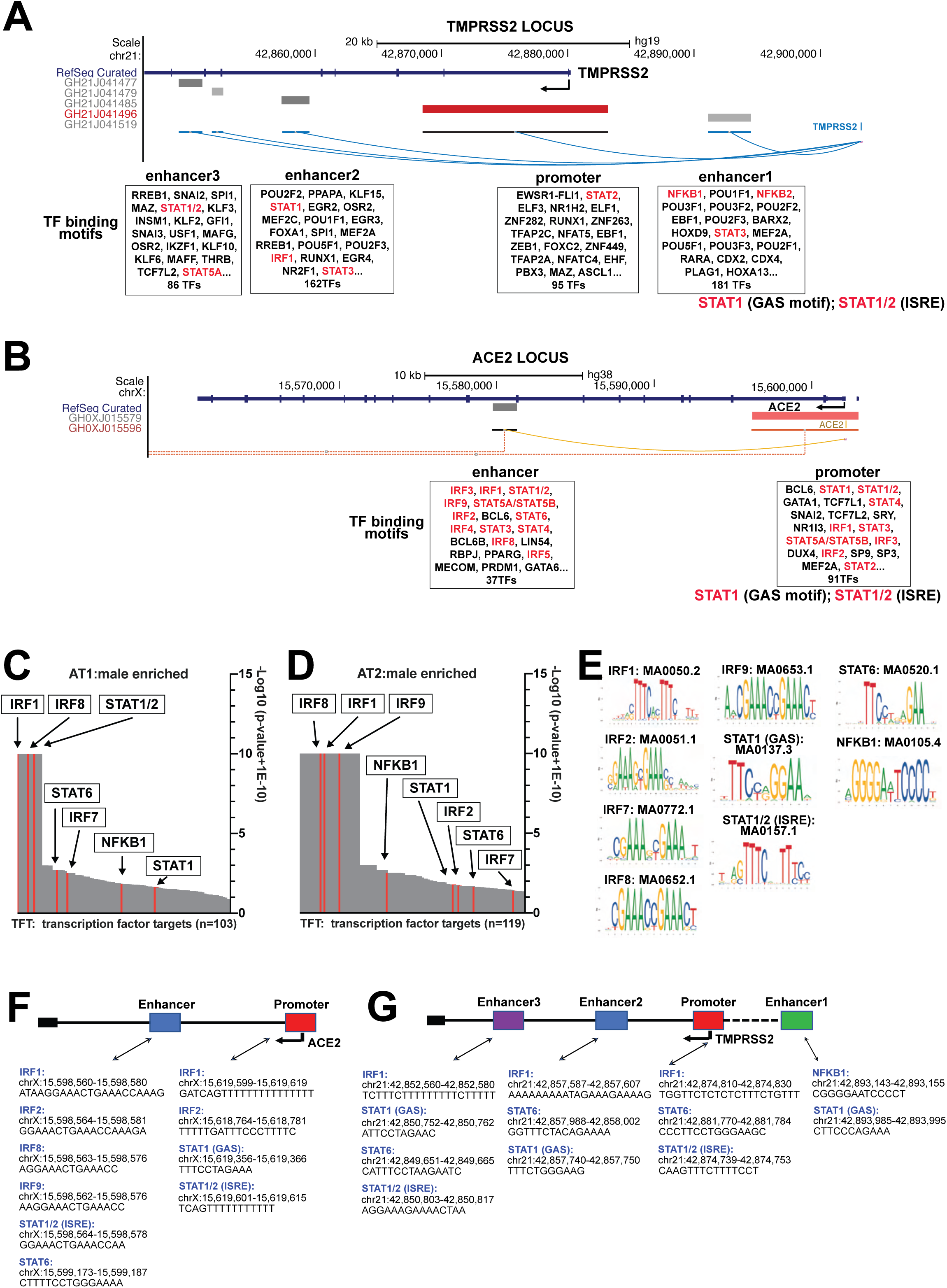
Lung-specific enhancers of TMPRSS2 are enriched in binding sites targeted by interferon transcription factors. (A) UCSC genome browser (http://genome.ucsc.edu) view of the *TMPRSS2* gene, with aligned transcription factors according to the binding motif from JASPAR (http://jaspar.genereg.net), and the regulatory elements and gene interactions from GeneHancer (https://genome.ucsc.edu/cgi-bin/hgTrackUi?db=hg19&g=geneHancer). (B) UCSC genome browser (http://genome.ucsc.edu) view of the *ACE2* gene, with aligned transcription factors according to the binding motif from JASPAR, and the regulatory elements and gene interactions from GeneHancer. (C) Male-specific enriched gene sets in “all transcription factor targets” from MSigDB database (http://www.gsea-msigdb.org/gsea/msigdb/index.jsp) for AT1 cells. The y-axis represents the value of log10-converted adjusted p-value. The x-axis displays the transcription factors corresponding to the gene sets. (D) Male-specific enriched gene sets in “all transcription factor targets” from MSigDB database for AT2 cells. The y-axis represents the value of log10-converted adjusted p-value. The x-axis displays the transcription factors corresponding to the gene sets. (E) Motifs of transcription factors involved in interferon signaling enriched in male AT1/AT2 cells. The motif logos are downloaded from JASPAR 2020 website. (F) Genomic location and sequence of the representative transcription factor binding sites in the promoter and enhancer regions of *ACE2*. (G) Genomic location and sequence of the representative transcription factor binding sites in the promoter and enhancer regions of *TMPRSS2*.

To functionalize this information, we conducted Gene Set Enrichment Analysis (GSEA) of the TF signatures enriched in male as compared to female AT1 and AT2 cells and identified several sexually dimorphic signatures expressed in AT1 and AT2 cells (Fig. 3C and 3D). Remarkably, STAT1, STAT1/2, various Interferon Response Factors (IRFs), and NF-*κ*B were the only TFs that coordinated a gene expression program enriched in male alveolar epithelium and had binding sites in the promoter and enhancers of *TMPRSS2* and *ACE2* (Fig. 3E-G). In contrast, although we did not identify statistically significant differences in the expression of hormone receptors (*AR*, *ESR1*, *ESR2*) between male and female AT1 and AT2 cells, we noted a trend toward gender divergent expression (fig. S3C and S3D). In addition, although canonical AR signatures were enriched in male AT1 and AT2 cells, the results did not reach statistical significance (fig. S3E). Finally, the ER_nongenomic_pathway signature was the only ER-related signature enriched in female AT1 or AT2 cells (fig. S3F). These results suggest that IFN and NF-*κ*B signaling are preferentially activated in normal male AT1 and AT2 cells *in vivo* as compared to their female counterpart; in contrast, AR signaling is inactive or weakly activated in these cells.

To corroborate these findings, we examined the enrichment of signatures from the Hallmark_gene_set, the Canonical_pathway_gene_set, and the GO_BP_gene_set in male versus female AT1 and AT2 cells. We found that several immune and inflammatory signatures are selectively enriched in male AT1 and AT2 cells. Importantly, signatures reflective of heightened IFN signaling featured amongst the top upregulated in male AT1 and AT2 cells as compared to their female counterparts (fig. S4A-F). Direct analysis of several signatures reflective of IFN signaling, including the HALLMARK_INTERFERON_ALPHA_RESPONSE signature, indicated that they are enriched in male AT1 and AT2 cells as compared to their female counterparts (Fig. 4A-D). Examination of the expression of several IFN signaling target genes confirmed their differential expression in the distal lung epithelium in the two genders (Fig. 4E, 4F, S4G, and S4H). Taken together, these findings suggest that sexually dimorphic IFN signaling controls the expression of SARS-CoV-2 entry receptors in the alveolar epithelium.

**Figure 4.**
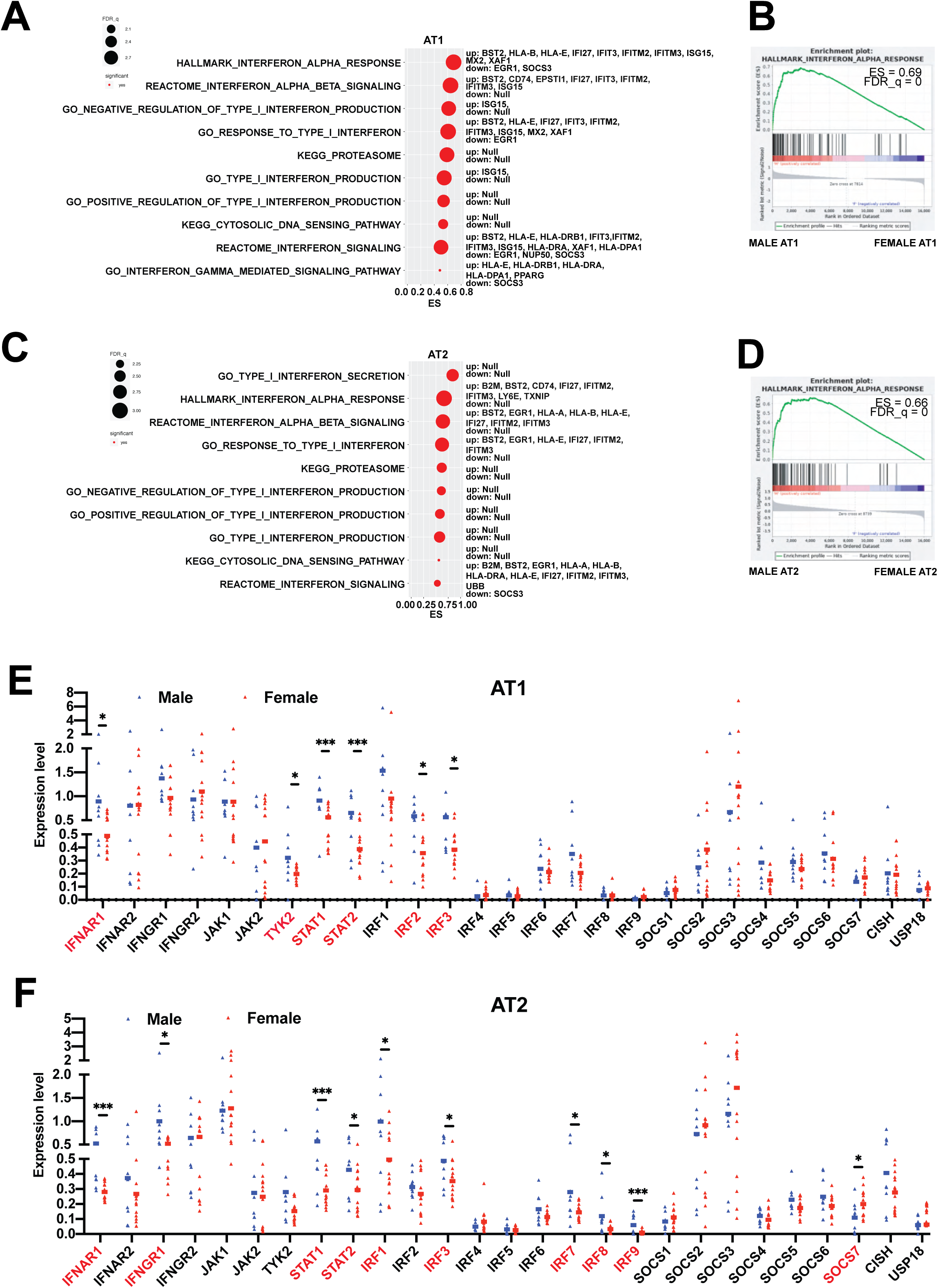
The activity of interferon signaling pathways is higher in male lung AT1 and AT2 cells than in female lung AT1 and AT2 cells. (A) Interferon-relevant signatures enriched for genes differentially expressed in AT1 female group versus AT1 male group using gene set enrichment analysis (GSEA). X-axis title “ES” represents the GSEA enrichment score. Y-axis represents the name of the signatures. Dot size represents the -log10 (FDR_q_value + 0.001). Dot color represents the significance (left); For each significant signature, the top differentially expressed genes, both up- and downregulated, are listed (right). (B) GSEA enrichment plot (score curves) of the top enriched gene set from Figure 2A. The y-axis represents enrichment score (ES) and the x-axis are genes (vertical black lines) represented in the gene set. The green line connects points of ES and genes. ES is the maximum deviation from zero as calculated for each gene going down the ranked gene list. The colored band at the bottom represents the degree of correlation of genes with the AT1 male group (red for positive and blue for negative correlation). Significance threshold set at FDR < 0.05. (C) Interferon-relevant signatures enriched for genes differentially expressed in AT2 female group versus AT2 male group using GSEA. X-axis title “ES” represents the GSEA enrichment score. Y-axis represents the name of the signatures. Dot size represents the -log10 (FDR_q_value + 0.001). Dot color represents the significance (left); For each significant signature, the top differentially expressed genes, both up- and downregulated, are listed (right). (D) GSEA enrichment plot (score curves) of the top enriched gene set from Figure 2C. The y-axis represents enrichment score (ES) and the x-axis are genes (vertical black lines) represented in the gene set. The green line connects points of ES and genes. ES is the maximum deviation from zero as calculated for each gene going down the ranked gene list. The colored band at the bottom represents the degree of correlation of genes with the AT2 male group (red for positive and blue for negative correlation). Significance threshold set at FDR < 0.05. (E) Expression of common interferon signaling genes in male and female AT1 cells. Genes showing significant difference between male and female group are highlighted in red. (F) Expression of common interferon signaling genes in male and female AT2 cells. Genes showing significant difference between male and female group are highlighted in red.

### Type I Interferon Signaling Upregulates the Expression of *ACE2* and *TMPRSS2* in Pulmonary Alveolar Epithelial Cells

To directly test the hypothesis that *ACE2* and *TMPRSS2* are IFN target genes in lung epithelium, we initially surveyed the expression of IFN receptors and JAK family kinases in primary human lung alveolar epithelial cells (AEpiC). qPCR indicated that these cells express IFNAR1, IFNAR2, IL-10Rβ, IFNGR1, IFNGR2, JAK1, TYK2, and lower levels of IFNLR1 and JAK2, suggesting that they can be stimulated by IFN-I (α, β), II (γ) and III (λ) (fig. S5A, S5B and S5C). Immunoblotting indicated that 20 nM IFNα promotes phosphorylation and accumulation of STAT1 and 2 in AEpiC cells, consistent with the finding that STAT1 and 2 are IFN target genes (47, 48). Tofacitinib, which preferentially inactivates JAK1 as compared to JAK2 and 3 and inhibits TYK2 much less efficiently (49), reversed both the phosphorylation and the accumulation of STAT1 and 2 (Fig. 5A). These results are consistent with the conclusion that IFNα-stimulated JAK-STAT signaling proceeds via the IFNAR1/2 heterodimer and associated JAK1/TYK2 kinases in AEpiC cells.

**Figure 5.**
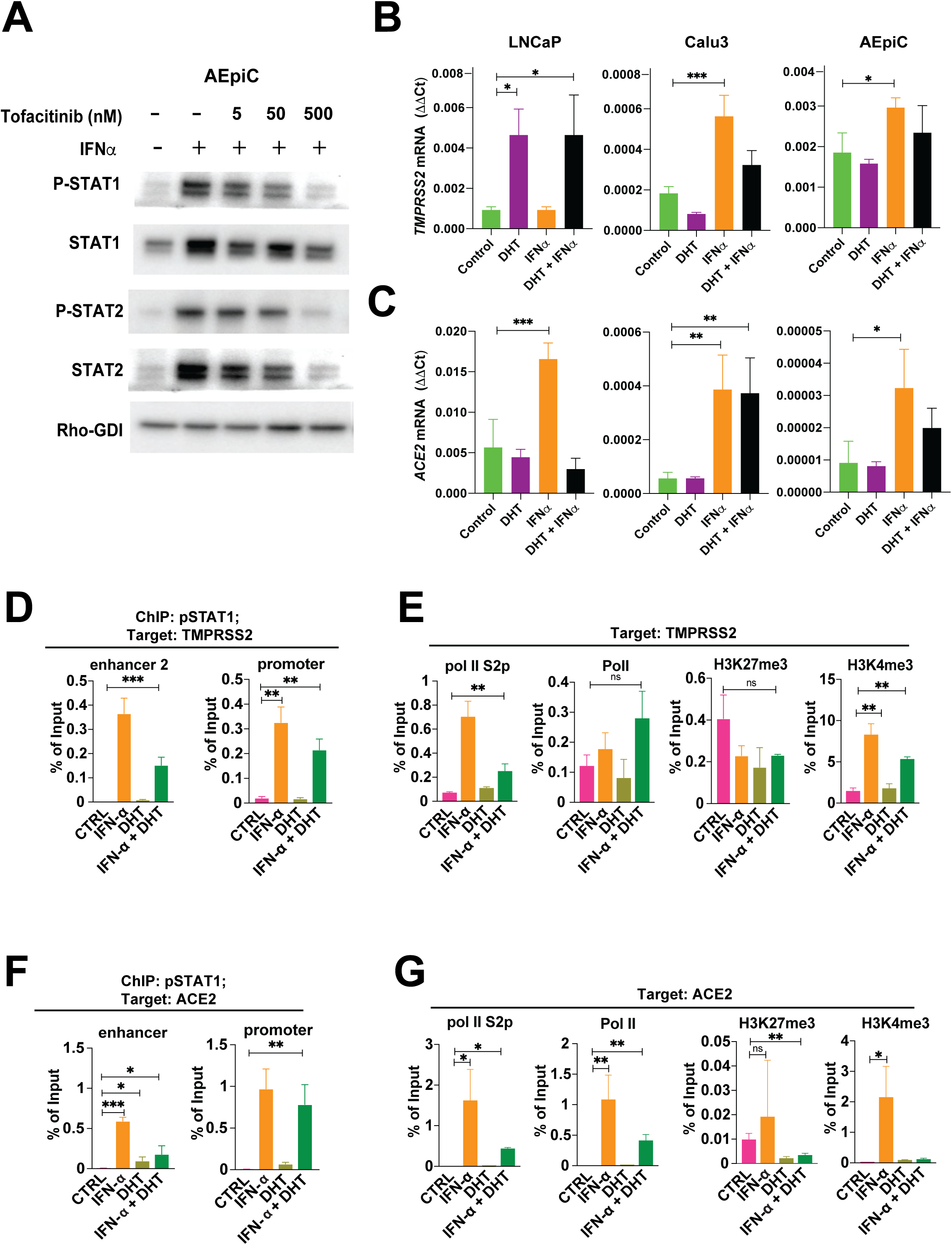
Interferon-stimulated JAK/STAT signaling, instead of AR-dependent signaling, transactivates the expression of SARS-CoV2 receptors, TMPRSS2 and ACE2, in human pulmonary alveolar epithelium. (A) To verify the activation of the downstream effectors in response to INF stimulation, AEpiC cells first pre-treated with JAK inhibitor, Tofacitinib, with indicated concentration for 1.5 hours, then cells were stimulated with 20 nM IFN⍺ for another 16 hours. Total lysates were lysed in ice-cold RIPA buffer and subjected to western blot to analyze the activation of STAT1 and STAT2, with GDI as an internal loading control. (B and C) Analysis of IFN⍺ and AR-dependent signaling in the expression of SARS-CoV-2 receptors in pulmonary alveolar epitheliums. LNCAP, Calu-3, and AEpiC cells were either treated with DHT alone (10 nM, 24 hours), IFN⍺ alone (20 nM, 16 hours), or in combination. After treatment, quantitative reverse transcription PCR (RT-qPCR) analysis of *TMPRSS2* and *ACE2* was performed to assess the expression of SARS-CoV-2 receptors, with *18S* ribosomal RNA as an endogenous control. (D and F) In order to directly examine whether p-STAT1 is capable of transactivates SARS-CoV-2 receptors in response to interferon or AR-dependent stimulation, 1×10^6^ AEpiC cells were either treated with DHT alone (10 nM, 24 hours), IFN⍺ alone (20 nM, 16 hours), or in combination. Cells were lysed in a non-denaturing lysis buffer and subjected to chromatin immunoprecipitation (ChIP) of p-STAT1, so as to investigate the enrichment of p-STAT1 on the enhancer and the promoter region of both *TMPRSS2(Top)* and *ACE2(Bottom)*. (E and G) The occupancy of both active transcription markers (H3K4me3, PoI II, and PoI II S2p) and a suppressive transcription marker (H3K27me3) on the regulatory region of SARS-CoV-2 receptors was also applied to investigate the transcriptional alteration of *TMPRSS2(Top)* and *ACE2(Bottom)* in copying with interferon response and AR-dependent signaling in AEpiC cells. Results are reported as mean ± SD. Comparisons between two groups were performed using an unpaired two-sided Student’s t test (p < 0.05 was considered significant). Comparison of multiple conditions was done with One-way or two-way ANOVA test. All experiments were reproduced at least three times, unless otherwise indicated.

To examine the capacity of IFN and androgen to regulate *ACE2* and *TMPRSS2* expression, we performed qPCR assays with LNCaP prostate adenocarcinoma cells, Calu-3 lung adenocarcinoma cells, and AEpiC cells, which were treated with IFNα(20 nM), androgen (10 nM DHT), or a combination of the two. Of note, the AEpiC cells consist of AT2 and, to a smaller extent, AT1 cells (50). The results indicated that DHT induces expression of *TMPRSS2* within 16 hours in LNCaP cells, as anticipated (Fig. 5B and 5C, left panel). However, DHT did not promote rapid expression of either *ACE2* or *TMPRSS2* in Calu-3 or AEpiC cells and treatment with the potent AR inhibitor enzalutamide (5 μM) did not reduce their level of expression in unstimulated cells (Fig. 5B and 5C, middle and right panels, and S5D). Notably, treatment with DHT did not enhance, but instead suppressed IFNα’s induction of *TMPRSS2* in Calu-3 and AEpiC. In contrast, IFNα induced expression of both *ACE2* and *TMPRSS2* in these two cell lines (Fig. 5B and 5C, middle and right panels). These results provide direct evidence that IFN signaling elevates the expression of SARS-CoV-2 entry receptors in lung alveolar epithelium. In addition, the lack of response to enzalutamide in AEpiC cells indicates that *TMPRSS2* is not an AR target gene in alveolar epithelial cells.

To examine the mechanism through which IFN signaling induces expression of *TMPRSS2*, we performed chromatin immunoprecipitation (ChIP)-qPCR assays using AEpiC cells treated with either IFN-α, DHT, or the combination. The results revealed that activated STAT1 (P-STAT1) binds to the promoter and enhancer 2 of *TMPRSS2* in IFNα-but not DHT-stimulated cells (Fig. 5D). This event was accompanied by a decrease of the suppressive mark H3K27me3, an increase of the activation mark H3K4me3, and a selective enrichment of activated Pol II (S2P-Pol II) at the promoter of *TMPRSS2* (Fig. 5E). Notably, treatment with DHT decreased the binding of P-STAT1 to the promoter and enhancer 2 of *TMPRSS2* and transcriptional activation of the gene in response to IFN (Fig. 5D,5E and S5E) and ChIP-qPCR assays indicated that this suppression did not involve direct binding of the AR to the lung-active enhancers or the promoter of *TMPRSS2* (fig. S5F). Therefore, IFN stimulation induces binding of activated STAT1 to the enhancer 2 and promoter of *TMPRSS2* and transcriptional activation of the gene. In contrast, DHT stimulation interferes with the IFN-stimulated expression of *TMPRSS2*.

By performing similar ChIP-qPCR assays, we found that P-STAT1 binds also to the promoter and intronic enhancer *ACE2* in IFNα-stimulated AEpiC cells and induces de-repression and transcriptional activation of the promoter (Fig. 5F and 5G). Interestingly, treatment with DHT decreased the binding of P-STAT1 to the promoter and enhancer of *ACE2* and transcriptional activation of the gene in response to IFN (Fig. 5F and G) as we had observed for *TMPRSS2*. However, ChIP-qPCR assays indicated that this suppression involved direct binding of the AR to the lung-active intronic enhancer and the promoter of *ACE2* (fig. S5G). These findings indicate that IFN signaling promotes the binding of P-STAT1 to the promoter and lung-specific enhancers of both *TMPRSS2* and *ACE2*, corroborating the hypothesis that sexually dimorphic IFN signaling leads to elevated levels of SARS-Cov-2 entry receptors in the male alveolar epithelium. In contrast, androgen does not exert these effects in lung epithelium.

### IFN-I and III Conspire to Upregulate the Expression of *ACE2* and *TMPRSS2* and Induce Robust Viral Entry

To identify the types of interferon that upregulates *ACE2* and *TMPRSS2* expression, we examined the effects of IFN-I (IFN⍺ and IFNβ), IFN-II (IFNɣ) and IFN-III (IFNλ) on the expression of *ACE2* and *TMPRSS2* in AEpiC cells. qPCR indicated that IFNβ and γ induce higher expression of *ACE2* as compared to IFNα and λ (Fig. 6A). In contrast, IFNλ induced higher expression of *TMPRSS2* as compared to IFNβ, and IFNα and γ proved ineffective (Fig. 6B). Immunofluorescent staining of non-permeabilized Calu-3 cells and qPCR analysis corroborated the preferential upregulation of ACE2 by IFNα and TMPRSS2 by IFNλ (Fig. 6C, S6A, and S6B). Immunoblotting analysis revealed that IFNα and IFNλ induce activation of JAK1 and TYK2 and, downstream, phosphorylation of STAT1 and 2 and formation of the ISGF3 complex (fig. S6C). As anticipated by JAK2’s exclusion from type I and type III IFN-Rs (51), IFNα and IFNλ did not induce its activation (fig. S6C, right). Consistent with their ability to engage molecularly distinct receptors, IFNα and IFNλinduced JAK-STAT signaling with divergent kinetics: rapid for IFNα and delayed for IFNλ (fig. S6C). Nevertheless, the two cytokines induced the accumulation of the proteins encoded by their target genes *STAT1* and *STAT2* with a similar kinetics (fig. S6C and S6D). Consistent with the low constitutive level of expression and the kinetics of accumulation of the *ACE2* mRNA in response to IFNα or λ (Fig. 6A), the canonical 75 kD membrane-anchored form of ACE2 was not detectable in untreated cells but was robustly induced at 12 hours and reached plateau expression at 24 hours. In contrast, TMPRSS2 was expressed under basal conditions but increased and reached plateau expression following the same kinetics as ACE2 (fig. S6C). Combinations of IFNαand IFNβ and triple combinations including IFNλ or γdid not stimulate expression of *ACE2* more than either IFNα or IFNβ alone (about 25-fold over control for both IFNαor IFNβ). Conversely, combinations of IFNλ with IFNαor IFNβ and triple combinations including IFNγ did not induce higher expression of *TMPRSS2* as compared to IFNλ alone (about 4-fold over control for IFNλ) (fig. S6E). In addition, treatment with JAK inhibitors such as fedratinib, ruxolitinib and tofacitinib, but not prednisone (corticosteroid), camostat or nafamostat (protease inhibitors) inhibited the IFNα dependent induction of ACE2 and TMPRSS2 (Fig. 6D). Together with the results of ChIP studies, these results provide strong evidence that *ACE2* and *TMPRSS2* are IFN target genes and that *ACE2* is predominantly controlled by IFN-I and *TMPRSS2* by IFN-III.

**Figure 6.**
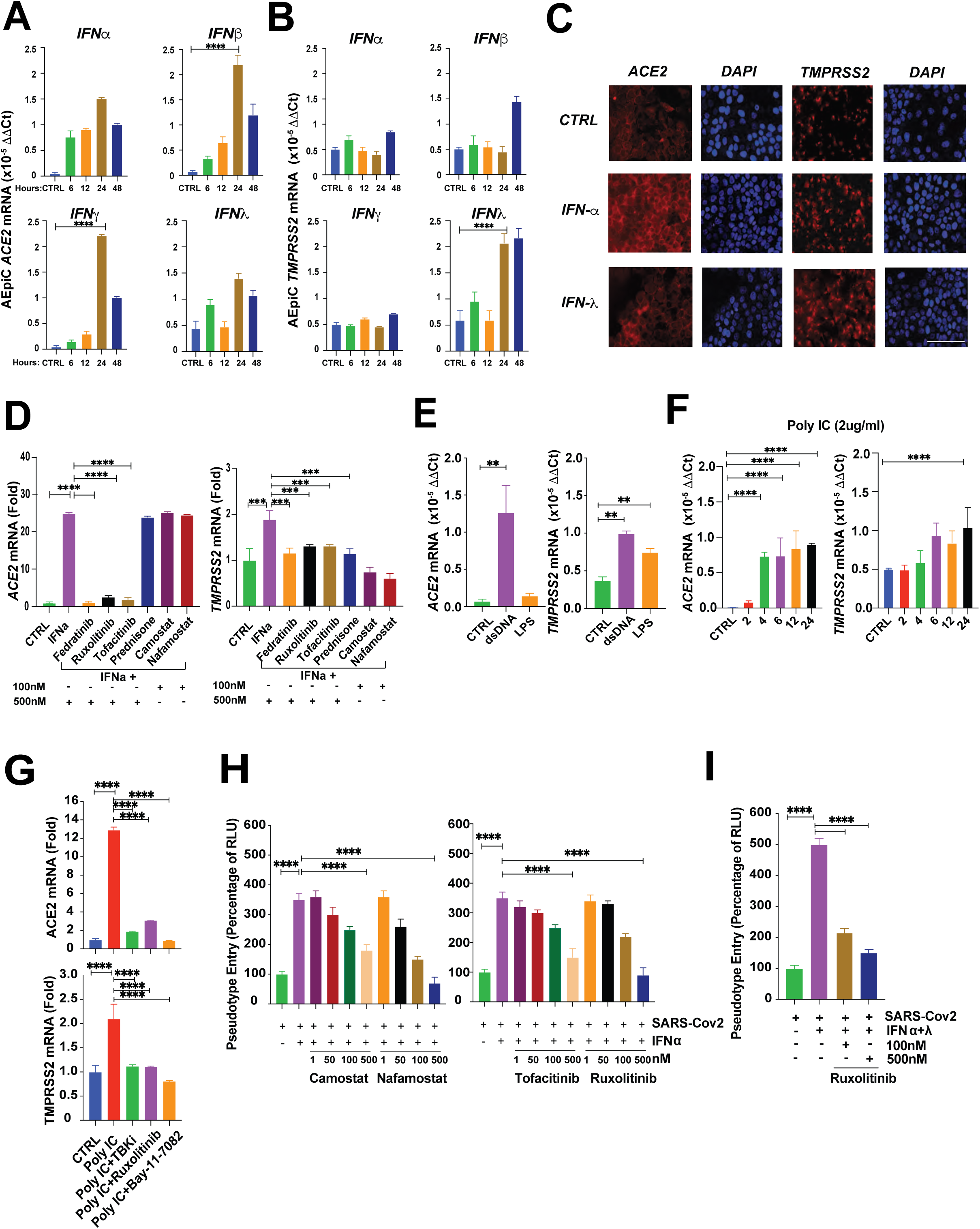
Interferons α/β and λ conspire to upregulate the expression of ACE2 and TMPRSS2 and induce robust viral entry. (A) *ACE2* mRNA expression level in response to different types of interferon stimulation. AEpiC cells treated with 10 nM IFN⍺ (upper left), 10 nM IFNβ (upper right), 10 ng/ml IFNɣ (lower left) or 1 µg/ml IFNλ (lower right) for the indicated times (6, 12, 24, 48 hours). Cells were subjected to RT-qPCR analysis to quantify the *ACE2* mRNA change, with *18S* as an endogenous control. (B) *TMPRSS2* mRNA expression level in response to different types of interferon stimulation. AEpiC cells treated with 10 nM IFN⍺ (upper left), 10 nM IFNβ (upper right), 10 ng/ml IFNɣ (lower left) or 1 µg/ml IFNλ (lower right) for the indicated times (6, 12, 24, 48 hours). Cells were subjected to RT-qPCR analysis to quantify the *TMPRSS2* mRNA change, with *18S* as an endogenous control. (C) Visualization of the alteration of SARS-CoV-2 receptors expression during type I and type III interferon stimulation. Calu-3 cells were either treat with PBS, 10 nM IFN⍺ (left) or 1 ug/ml IFNλ (right) for 16 hours. Then, cells were subjected to immunofluorescent stain to investigate the abundance of ACE2 and TMPRSS2. DAPI was used for nuclei staining (Scale bar = 100 um). (D) The Efficacy of different JAK and protease inhibitors in the blockade of SARS-CoV-2 pseudotype entry. AEpiC cells were incubated for 12 hours at different concentrations of fedratinib, ruxolitinib, tofacitinib, camostat, and nafamostat, with PBS or 10 nM IFN⍺ for 16 hours before viral infection. The transduction efficiency of the virus was quantified 48 hours post-transduction by measuring the activity of firefly luciferase in cell lysates. (E) AEpiC cells were transfected with 2 ug/ml dsDNA or 100 ng/ml LPS for 12 hours. *ACE2* (left) and *TMPRSS2* (right) mRNA expression measured with RT-qPCR, with *18S* as an endogenous control. (F) AEpiC cells were transfected with 2 ug/ml poly (I: C) for different time points as indicated. *ACE2* (left) and *TMPRSS2* (right) mRNA expression measured with RT-qPCR, with *18S* as an endogenous control. (G) AEpiC were transfected with 2 ug/ml poly (I: C) without or with 5 µM TBKi (GSK8612),500 nM ruxolitinib, or 1 µM BAY-11-7082. RT-qPCR was performed for ACE2 (top) and TMPRSS2 (bottom), with *18S* as an endogenous control. (H) JAK inhibitors block the Type I and Type III interferon-induced SARS-CoV-2 pseudotype entry. AEpiC were cells incubated for 12 hours with different concentrations of camostat, nafamostat (Left), fedratinib and ruxolitinib (Right) and were stimulated with either PBS or 10 nM IFN⍺ for 16 hours before viral infection. The transduction efficiency of the virus was quantified 48 hours post-transduction by measuring the activity of firefly luciferase in cell lysates. (I) AEpiC were cells incubated for 12 hours with different concentrations of ruxolitinib and were stimulated with either PBS or 10 nM IFN⍺ plus 1 ug/ml IFNλ for 16 hours before viral infection. The transduction efficiency of the virus was quantified 48 hours post-transduction by measuring the activity of firefly luciferase in cell lysates. Results are reported as mean ± SD. Comparisons between two groups were performed using an unpaired two-sided Student’s t test (p < 0.05 was considered significant). Comparison of multiple conditions was done with One-way or two-way ANOVA test. All experiments were reproduced at least three times, unless otherwise indicated.

IFN-I and III cooperate to restrict viral infection of epithelial cells, including those lining the airways of the lung. Newly infected cells produce IFNs in response to activation of pattern recognition receptors, including cGAS-STING (51, 52). To examine if activation of such pathways and, hence endogenous production of IFNs, can upregulate expression of *ACE2* and *TMPRSS2* in alveolar epithelial cells, we initially transduced dsDNA or LPS into AEpiC cells. qPCR revealed that dsDNA induces expression of both entry receptors, but LPS only upregulates TMPRSS2 (Fig. 6E). Since LPS is recognized by the Toll-like Receptor 4 (TLR4), which predominantly impinges on NF-*κ*B signaling, the latter observation implies that TMPRSS2 may be induced also by NF-*κ*B (53). To better model the effect of the viral RNA of SARS-CoV-2, we used Poly IC. As shown in Figure 6F, poly IC rapidly induced expression of *ACE2* and *TMPRSS2*, suggesting that initial viral entry stimulates expression of viral entry receptors, potentially facilitating the entry of additional viruses. The expression of *ACE2* and *TMPRSS2* induced by Poly IC was suppressed by treatment of the cells with an inhibitor of TBK kinase (GSK8612), which controls expression of IFNs via NF-*κ*B (54), the JAK1/2 and TYK2 inhibitor ruxolitinib (49) and the NF-*κ*B inhibitor BAY-11-70-82 (55)(Fig. 6G). This pattern of inhibition is consistent with the signaling mechanisms that enable cGAS-STING to induce the expression of IFNs. Together, these observations suggest that SARS-CoV2 exploits the ability of its RNA to induce the production of IFN to radically increase the expression of its entry receptors on alveolar epithelial cells.

To determine if IFN can enhance the entry of SARS-CoV2 into alveolar epithelium, we used replication-defective lentiviral particles bearing coronavirus S proteins. Pseudotyped viral particles have been shown to enter into cells by binding to ACE2 and TMPRSS2 (9). Firstly, we tested if knocking down ACE2 and TMPRSS2 could hinder the entry of SARS-2S pseudotyped lentivirus in CALU3 cells. As show in Figure S6F and S6G, reduced expression of either one of the receptors significantly reduced the viral entry. We then stimulated AEpiC cells with IFN⍺ for 16 hours and infected them with LVM-SARS-CoV-2_S, a SARS-2S pseudotyped lentivirus encoding luciferase. Stimulation with IFNα increased viral entry by 3-fold, and pre-treatment with the TMPRSS2 protease inhibitors camostat and nafamostat inhibited it, confirming the dependency of this process on S protein binding and activation by TMPRSS2 (Fig. 6H). Pre-treatment with the JAK inhibitors tofacitinib and ruxolitinib also suppressed the entry of the virus induced by IFN (Fig. 6H). To model the microenvironment of the early infection of alveolar epithelium, we repeated the experiment by exposing the AEpiC cells to a mixture of IFNα and IFNλ. In agreement with its capacity to induce maximal expression of both ACE2 and TMPRSS2, the combination promoted a 5-fold increase in viral entry, which was largely reversed by ruxolitinib (Fig. 6I). These findings suggest that IFN-I and IFN-III can substantially upregulate the expression of viral entry receptors during the early phase of infection of the alveolar epithelium and that JAK inhibitors can interfere with this process.

### TNF⍺ Models Severe SARS-COV2 Disease and Cytokine Storm in Pulmonary Alveolar Epithelial Cells

To model the effect of cytokines present in the lung of COVID-19 patients affected by advanced lung disease, we tested IFNγ, TNFα, and additional cytokines, which have been found to correlate with or participate in disease progression (56–59). Notably, TNFα induced expression of *TMPRSS2* but not *ACE2*, whereas IFNγ induced expression of *ACE2* but not *TMPRSS2*. The two cytokines in combination did not exert a higher effect as compared to either one singly (Fig. 7A and S7A). None of the 14 additional cytokines tested induced a significant increase in the expression of *ACE2* or *TMPRSS2* (fig. S7B). Whereas the IFNγ receptor signals via JAK1/2 and STAT3, the TNFα receptor signals predominantly through activation of NF-*κ*B (51, 60). Accordingly, ChIP experiments revealed that IFNɣ promotes the binding of p-STAT3 to the enhancer and promoter of *ACE2* but not of *TMPRSS2* (Fig. 7B and S7C). In contrast, TNF⍺-promotes binding of the NF-*κ*B-p65 complex to the enhancer and promoter of *TMPRSS2* but not of *ACE2* (Fig. 7C and S7C). These findings indicate that at concentrations inferior to those inducing inflammatory cell death (56), TNF⍺ and IFNγ induce a robust upregulation of SARS-CoV-2 entry receptors in the alveolar epithelium.

**Figure 7.**
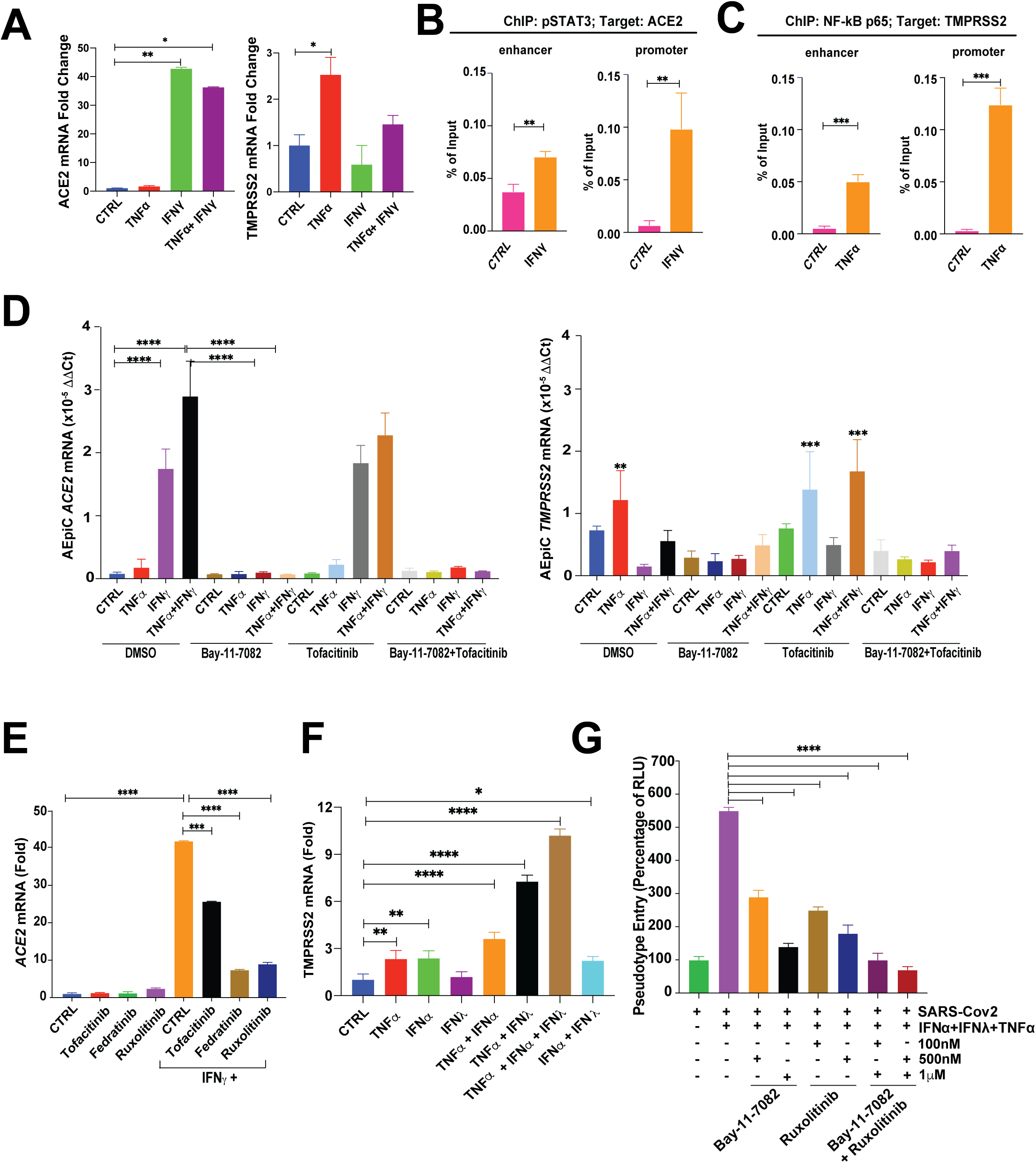
TNF⍺ models severe SARS-COV2 disease and cytokine storm in pulmonary alveolar epithelial cells. (A) SARS-CoV-2 receptors mRNA expression level in response to TNF⍺, IFNɣ, cytokines presented in the advanced lung disease of COVID-19 patients. AEpiC cells were treated with PBS, 10 ng/ml TNF⍺, 10 ng/ml IFNɣ, or in combination for 24 hours. *ACE2* (left) and *TMPRSS2* (right) mRNA expression measured with RT-qPCR, with *18S* as an endogenous control. (B) Enrichment of p-STAT3 on the enhancer and the promoter of *ACE2* from ChIP-qPCR in control and IFNɣ of AEpiC cells. (C)Enrichment of NF-kB p65 on the enhancer and the promoter of *TMPRSS2* gene from ChIP-qPCR in control and TNF⍺ of AEpiC cells. (D) The efficacy of NF-kB inhibitor and JAK1 inhibitor in the suppression of SARS-CoV-2 receptors expression. AEpiC cells were pre-treated with either 1 µM BAY-11-7082, 500 nM tofacitinib, or in combination for 12 hours and then stimulated with either 10 ng/ml TNF⍺, 10 nM IFN⍺, 10 ng/ml IFNɣ, or in combination for another 24 hours. RT-qPCR analysis was performed to assess the mRNA expression level of *ACE2* (top) and *TMPRSS2* (bottom), with *18S* as an endogenous control. (E) The dependency of different JAKs on the *ACE2* expression during IFNɣ stimulation. AEpiC cells were pre-treated with either 500 nM tofacitinib, 500 nM fedratinib, or 500 nM ruxolitinib for 12 hours and then stimulated with 10 ng/ml IFNɣ for another 24 hours. RT-qPCR analysis was performed to assess the mRNA expression level of *ACE2* (top) and *TMPRSS2* (bottom), with *18S* as an endogenous control. (F) AEpiC cells were treated with either PBS, 10 ng/ml TNF⍺, 10 nM IFN⍺, 10 ng/ml IFNλ, or in combinations for 24 hours. *TMPRSS2* mRNA expression measured by RT-qPCR, with *18S* as an endogenous control. (G) The efficacy of NF-kB inhibitor and JAK1/2 inhibitor in the blockade of SARS-CoV-2 pseudotype entry in the COVID-19 advanced lung disease model. AEpiC cells were incubated for 12 hours with different concentrations of BAY-11-7082, ruxolitinib, or in combination, and stimulated without or with 10 nM IFN⍺, 1 ug/ml IFNλ, and 10 ng/ml TNF⍺ for 16 hours before viral infection. The transduction efficiency of the virus was quantified 48 hours post-transduction by measuring the activity of firefly luciferase in cell lysates. Results are reported as mean ± SD. Comparisons between two groups were performed using an unpaired two-sided Student’s t test (p < 0.05 was considered significant). Comparison of multiple conditions was done with One-way or two-way ANOVA test. All experiments were reproduced at least three times, unless otherwise indicated. Supplementary Figure Legends

We next asked if inhibition of JAK-STAT and NF-*κ*B signaling would suppress the expression of *ACE2* and *TMPRSS2* induced by a combination of IFNγ and TNFα. Notably, BAY-11-7082 suppressed the expression of *ACE2* induced by IFNγ or IFNγ and TNFα, but the JAK1 inhibitor tofacitinib did not (Fig. 7D). While the inhibition of the effect of IFNγ by BAY-11-7082 may arise from the crosstalk between IRF, NF-*κ*B, and JAK-STAT pathways (61), the inability of tofacitinib to block the effect of IFNγ was unexpected. We therefore tested optimal concentrations of tofacitinib (JAK1i), fedratinib (JAK2i), and ruxolitinib (JAK1/2i) and found that only the latter two compounds suppress the upregulation of ACE2 induced by IFNγ (Fig. 7E). These results suggest that the upregulation of *ACE2* induced by IFNγ depends more on JAK2 than JAK1 under our experimental conditions. As anticipated, BAY-11-7082 suppressed the expression of *TMPRSS2* induced by TNFα and tofacitinib inhibited that induced by IFNγ (Fig. 7D). These results suggest that combined inhibition of JAK-STAT and NF-*κ*B signaling may inhibit the expression of SARS-CoV-2 entry receptors induced by TNFα and IFNγ during the cytokine storm that typifies advanced disease.

We finally reasoned that IFN⍺ and λ would continue to be produced during advanced disease as long as new alveolar epithelial cells are infected. We therefore asked if a combination of IFN⍺ and λ and TNFα could promote a simultaneous and substantial upregulation of both ACE2 and TMPRSS2. Notably, we found that the triple combination induces higher expression of both *TMPRSS2* and *ACE2* as compared to all double combinations or each cytokine alone (Fig. 7F). Moreover, the triple combination was superior to the combination of IFNγ and TNFα (Fig. 7A and 7F). Intriguingly, the NF-*κ*B inhibitor BAY-11-7082 or the JAK inhibitor ruxolitinib substantially inhibited the entry of SARS-2S pseudotyped lentivirus into AEpiC cells stimulated with IFN⍺ and λ and TNFα. When used in combination, the two inhibitors completely suppressed viral entry (Fig. 7G). These results demonstrate that IFNs and TNF⍺ substantially upregulate ACE2 and TMPRSS2 during severe disease and the combined inhibition of their signaling pathways may ameliorate disease progression.

## Discussion

In this study, we provide evidence that sexually dimorphic IFN and, possibly, NF-*κ*B signaling upregulates expression of the viral entry receptors ACE2 and TMPRSS2 in male alveolar type I and II cells, potentially explaining why progression to SARS occurs much more frequently in this gender. Examination of primary alveolar epithelial cells indicates that IFN-I and III, which are involved in the protective immunity of epithelial surfaces (62, 63), dramatically upregulate the expression of *ACE2* and *TMPRSS2* and facilitate viral entry of an S protein-pseudotyped virus. TNF-α and IFNγ, which are produced during the cytokine storm associated with the lethal phase of the disease (64), upregulate the entry receptors by a similar amplitude. Finally, pharmacological inhibition of JAK1/2 or both JAK1/2 and NF-*κ*B suppressed viral entry of the pseudotyped virus in cells stimulated with optimal combinations of IFNs or with TNFα and IFNγ, respectively. In addition to providing a potential explanation for the predominance of severe disease in men, these results indicate that SARS-Cov-2 hijacks viral immunity mechanisms to facilitate viral spreading and suggests novel and distinct therapeutic strategies for the prevention and treatment of SARS.

Recent studies using long-term human distal lung organoids have consolidated the view that SARS-CoV-2 can productively infect AT2 cells, resulting in an innate immune response, cell death and downregulation of surfactant expression (16, 65, 66). Upon examining a combined single-cell RNA-seq dataset consisting of 93,770 transcriptomes from 24 normal individuals, we found that *ACE2* is expressed in only a very small proportion of AT1 and, as previously reported (11, 15), AT2 cells (<1%). In contrast, *TMPRSS2* is expressed in a large fraction of both cell types (ca. 36%). Since approximately 50% of ACE2+ cells also express TMPRSS2, double-positive AT1 and AT2 cells are extremely rare in the normal lung (0.22% and 0.44%, respectively). We thus posited that additional signaling mechanisms upregulate the expression of *ACE2* and *TMPRSS2* in these cells and thereby facilitate initial viral entry as well as the progression of the infection, especially in the face of protective innate immunity mechanisms.

The mechanisms underlying the male prevalence of SARS are poorly understood. Intriguingly, we found that the expression of *ACE2* and *TMPRSS2* is significantly higher in normal alveolar epithelial cells from males as compared to females. Previous studies have shown that women mount more robust immune responses against viruses and vaccines and exhibit superior immune-mediated tissue repair as compared to males (33). In addition, clinical studies have shown that male patients with moderate COVID-19 have defective T cell responses that correlate with disease severity (30). While it is likely that differences in immune responses between the genders contribute to the higher disease severity in males, our finding that males possess about twice as many ACE2+ and TMPRSS2+ AT1 and AT2 cells as compared to females suggests that the male alveolar epithelium is more prone to SARS-CoV-2 infection because it contains a larger number of cells co-expressing the viral entry receptor and co-receptor.

Analysis of the ENCODE database identified distal enhancers of *ACE2* and *TMPRSS2* active in lung tissue. In contrast, the canonical AR-regulated enhancer of *TMPRSS2*, which is active in prostate epithelial cells, is located proximally and was not active in the lung. Transcription factor binding motif and GSEA suggested that several JAK-activated transcription factors, including STAT1 and 2, and IFN-induced transcription factors, including IRF1, are induced and bind to these novel enhancers and the promoter of *TMPRSS2* and *ACE2* to a larger extent in male AT1 and AT2 cells as compared to their female counterparts. In contrast, we did not detect a statistically significant difference in the level of expression or activity of AR between male and female AT1 and AT2 cells. Provocatively, GSEA revealed that several IFN-regulated signatures are amongst the top upregulated in male AT1 and AT2 cells as compared to their female counterparts. Consistently, several canonical IFN target genes were expressed at higher levels in male AT1 and AT2 cells as compared to their female counterpart. These results suggest that both *ACE2* and *TMPRSS2* are IFN target genes in alveolar epithelial cells and that sexually dimorphic IFN signaling leads to higher levels of expression of both viral entry receptors in males, potentially explaining the increased susceptibility of males to SARS.

To directly confirm these findings, we examined the effect of IFN signaling on the expression of *ACE2* and *TMPRSS2* in primary human lung alveolar epithelial cells. We found that these cells respond to stimulation with IFNαby activating STAT1 and STAT2 in a JAK-dependent manner. Consistent with the notion that STAT1 and 2 are not only transcriptional activators but also target genes in the IFN signaling pathway (47, 48), IFNα robustly activated their expression, providing a feed-forward mechanism for amplification of signaling in the alveolar epithelium. Notably, treatment of human lung alveolar epithelial cells with IFN⍺ promoted binding of p-STAT1 to the promoter and the enhancer of *ACE2*, transcriptional activation of the gene, and expression of ACE2. Similarly, IFNα induced binding of p-STAT1 to the promoter and the lung-active distal enhancer of *TMPRSS2*, transcriptional activation of the gene, and expression of TMPRSS2. In contrast, treatment with androgen did not stimulate expression of the two entry receptors; in fact, it interfered with IFN-induced p-STAT1 binding to the regulatory elements of *ACE2* and *TMPRSS2* and therefore reduced their expression. Moreover, we found that the second-generation AR inhibitor enzalutamide does not reduce the expression of *ACE2* and *TMPRSS2* in primary alveolar epithelial cells. In fact, enzalutamide increased the expression of TMPRSS2 in these cells, presumably by interfering with the ability of autocrine IFN signaling to induce STAT1/2 binding to the promoter and distal enhancer of *TMPRSS2*. Together, our results demonstrate that IFN signaling upregulates the expression of viral entry receptors in primary alveolar epithelial cells. Notably, these conclusions are not consistent with the previously held view that androgen contributes to the regulation of *TMPRSS2* in the distal lung, which was based on indirect evidence (21), retrospective clinical associations (22), or mechanistic studies in prostate cancer LNCaP cells (67).

Although alveolar epithelial cells express type I, II, and III IFN-Rs, IFNα and β induce expression of *ACE2* to a much larger extent (>20 fold over control) as compared to other IFNs. In contrast, IFNλ is the most potent inducer of *TMPRSS2* (>7 fold over control). Since the basal expression of *ACE2* is much lower than that of *TMPRSS2*, co-stimulation with type I and II IFN induces a relative overexpression of *TMPRSS2* as compared to *ACE2*, suggesting that multiple TMPRSS2 co-receptors may assist a single ACE2 receptor in mediating viral entry in alveolar epithelial cells. It is hypothesized that initial viral entry into naïve epithelial cells induces expression of both type I and III IFNs through pattern recognition receptors (62, 63). Intriguingly, we found that transduction of Poly IC, mimicking viral RNA, robustly upregulates the expression of *ACE2* and *TMPRSS2* in alveolar epithelial cells, suggesting that viral entry and the ensuing liberation of viral RNA into the cytoplasm can induce production of IFNs and upregulation of viral entry receptors in a feed-forward mechanism. Importantly, our results also suggest that secreted type I and III IFN will upregulate the expression of viral entry receptors in adjacent epithelial cells, facilitating their infection. Consistent with the role of JAK-STAT signaling in this process, type I and III IFN stimulated entry of an S protein-pseudotyped virus in alveolar epithelial cells and JAK inhibitors reversed this process. These results suggest that JAK inhibitors may be effective in preventing the transition of COVID-19 disease to SARS in addition to ameliorating disease progression in SARS patients, as suggested by recent clinical trials (68, 69).

The cytokine shock syndrome often underlies the progression of SARS-CoV-2 to the lethal stage (64). We found that TNF⍺ upregulates the expression of TMPRSS2, whereas INFγ controls the expression of ACE2. In contrast, 14 other cytokines potentially involved in the cytokine storm do not affect the expression of either one of the two viral entry receptors. Consistent with the prominent role of NF-*κ*B in TNFα receptor signaling and JAK-STAT3 in IFNγ signaling (70, 71), a combination of the IKK/NF-*κ*B inhibitor BAY-11-7082 and the JAK inhibitor ruxolitinib completely suppressed viral entry induced by the combination of TNF⍺ and INFγ in alveolar epithelial cells. Intriguingly, prior studies have indicated that the combination of TNF⍺ and INFγ can induce PANoptosis (inflammatory cell death) of epithelial cells and promotes death in mice infected with SARS-CoV2 (56). Our results suggest that the ability of TNF⍺ and INFγ to upregulate entry receptors and promote further spreading of the virus in infected cells may potentiate the toxic effect of the cytokine storm on lung epithelial cells. Irrespective of the relative contribution of viral spreading and cytokine toxicity to disease progression, our results suggest that combinations of JAK inhibitors and clinically approved NF-*κ*B inhibitors, such as salicylates (72), should be tested in the advanced stage of COVID-19.

Recent studies have shown that even single virions can productively infect AT2 cells in alveolar organoids. Notably, whereas high levels of IFN limit further infection, resulting in modest viral burden, low levels of IFN exerts the opposite effect, suggesting that IFN signaling can exert a bimodal effect depending on its dose (16, 66). We propose that viral mechanisms enable newly infected cells to titrate the production of type I and III IFNs to a level that is insufficient to mediate viral restriction but is sufficient to upregulate viral entry receptors. Consistently, it has been observed that SARS-CoV-2 infection drives lower antiviral transcription marked by low IFN-I and IFN-III levels as compared to common respiratory viruses (73). By necessity, most of the studies on viral proteins suppressing IFN signaling have been conducted by using overexpression of individual viral genes and, thus, cannot provide definitive information on the level of IFN signaling induced in naturally infected cells (74–78). Future studies using a recently developed *trans*-complementation system will contribute to a better understanding of how SARS-CoV-2 fine-tunes IFN signaling to facilitate its spreading in the host (79). We further propose that type I IFN immunity plays a similar bimodal role also in late-stage disease. In fact, it has been reported that inborn errors of IFN-I immunity or autoantibodies to IFN-I are present, at a low rate, in patients with a life-threatening disease, but they are completely absent in those with mild disease (80, 81). In contrast, immunophenotyping of patients suggest that that the IFN-I response exacerbate inflammation in severe COVID-19 (28) and IFN-I and III disrupt lung epithelial repair during recovery from viral infection (82).

In conclusion, our study indicates that sexually dimorphic IFN-I and III signaling upregulates the expression of the SARS-CoV-2 entry receptors *ACE2* and *TMPRSS2* in lung alveolar epithelium, providing a potential mechanism for the male-biased incidence of SARS in COVID-19 patients. We also demonstrate that SARS-CoV-2 hijacks IFN-I and III signaling to upregulate the viral entry receptors and hence facilitate viral spreading during initial infection of the alveolar epithelium. Furthermore, induced as components of the cytokine storm, TNF-α and IFN-II produce a similar upregulation of viral entry receptors and a direct cytopathic effect during the late phase of the disease. Based on these results, we suggest that JAK1/TYK2 inhibitors, such as ruxolitinib, may be used in conjunction with antivirals to halt viral spread from the upper respiratory tract to the distal lung, reducing the incidence of SARS. Furthermore, combinations of JAK1/2, such as baricitinib, and NF-*κ*B inhibitors, such as salicylate, may be more efficacious as compared to current treatments during advanced SARS in COVID-19. In addition to informing our ongoing understanding of COVID-19 pathophysiology, these findings suggest novel therapeutic strategies and regimens for the prevention and treatment of COVID-19 SARS.

We were unable to include several scRNAseq datasets in this study due to unavailability of sex information. Our results on sexually dimorphic IFN signaling and expression of ACE2 and TMPRSS2 in alveolar epithelial cells are therefore based on a comparison of a relatively small number of individual samples. In addition, we did not validate these results by using an independent approach. Finally, although our results suggest that SARS-Cov-2 does not suppress IFN signaling in infected cells as profoundly as other viruses, this model will need to be validated by using the wild-type virus or a trans-complementation system.

## Methods

### Study Design

We have used public scRNAseq datasets to compare the level of expression of SARS-CoV-2 entry receptors in male and female lung alveolar epithelial cells. By using transcription factor binding motif analysis and gene set enrichment analysis, we have then identified the signaling pathways potentially able to regulate the expression of TMPRSS2 and ACE2 in alveolar type I and II cells. To confirm these observations, we have examined the ability of IFN and NF*κ*B signaling to elevate the expression of both entry receptors in primary alveolar epithelial cells by using ChIP-Q-PCR, Q-PCR, and immunoblotting with antibodies to activated JAK and STAT isoforms. Finally, we have used a pseudo-typed lentiviral reporter vector to study the effect of IFN and NF*κ*B signaling and specific inhibitors on viral entry in primary alveolar epithelial cells.

### Reagents

A list of reagents, including antibodies, probes, cell lines, chemical reagents and software, can be found in the Resources table (Supplementary Table 1).

### Single cell RNA sequencing datasets

Three publicly available scRNA-seq datasets were obtained as follows: 1) processed data including count and metadata tables of healthy lung tissue was downloaded from Figshare (https://doi.org/10.6084/m9.figshare.11981034.v1); 2) h5 files of normal lungs were extracted from the Gene Expression Omnibus (GEO) database (https://www.ncbi.nlm.nih.gov/geo/) under accession number GSE122960; and 3) processed data including count and metadata tables of human lung tissue was acquired from GSE130148. All three datasets were generated on Illumina HiSeq 4000. Characteristics of all the samples containing sample name, sex, age and smoking status are provided in Supplementary Table 2.

### Single cell RNA sequencing data analysis

Count matrix was used to create a Seurat object for each dataset, and the three Seurat objects were further merged into a new Seurat object with the resulting combined count matrix. The merged matrix was first normalized using a global-scaling normalization method “LogNormalize” in Seurat v.3.2.0 with default parameters. To detect the most variable genes used for principal component analysis (PCA), variable gene selection was performed and the top 2,000 variable genes were then selected using the ‘vst’ selection method in Seurat FindVariableFeatures function. All genes were scaled in the scaling step and PCA was performed using the selected top 2,000 informative genes. To do batch effect correction, Harmony algorithm was run on the top 50 PCA embeddings using RunHarmony function. Then UMAP calculation was performed on the top 30 Harmony embeddings for visualizing the cells. Meanwhile, graph-based clustering was done on the harmony-reduced data. The final resolution was set to 0.2 to obtain a better clustering result.

### Generation of histone modification enrichment profiles and transcription factor binding motif analysis

H3K27ac and H3K4me3 binding profiles were constructed using publicly available ENCODE datasets. Accession numbers for all ENCODE datasets used can be found in Encode Data Sets Table (Supplementary Table 3). The data were visualized with UCSC genome browser (83, 84). Predicted enhancer regions of TMPRSS2 were identified using the GeneHancer tool within Track Data Hubs of UCSC genome browser (85, 86). Promoter and enhancer associated transcription factors were predicted by JASPAR (87).

### Gene set enrichment analysis

Gene set enrichment analyses (GSEA) were performed according to the instructions. Gene sets of Hallmark Collection, Canonical Pathway (including KEGG Pathway, Biocarta Pathway, Reactome Pathway and PID Pathway), and GO Biological Process were used. All transcription factor targets are from the Molecular Signatures Database (MSigDB) version 7.1. The interferon signatures were searched from MSigDB with the Keyword: interferon and the Search Filters: “H: hallmark gene sets; --CP: canonical pathways; --GO: Gene Ontology”.

### Cell lines and cell culture

All cells were incubated at 37°C and 5% CO_2_. AEpiC cells were obtained from Cell Biologics (#H-6053), and Calu-3 cells were obtained from ATCC (#). AEpiC cells were incubated with Alveolar Epithelial Cell Medium (ScienCell, #3201) supplemented with 2% fetal bovine serum, penicillin, streptomycin, and epithelial growth supplement. Calu-3 cells were incubated with Dulbecco’s Modified Eagle Medium supplemented with 10% fetal bovine serum, penicillin, and streptomycin.

### Western blotting

For immunoblotting, cells were washed once with PBS and lysed in either RIPA buffer with protease inhibitor or Laemmli sample buffer with BME. Samples were quantified with Piece BCA Protein Assay Kit (Thermo Scientific, #23225) where applicable, and boiled for 5 minutes before gel loading. Lysates were run on 4-15% precast electrophoresis gels, and transferred to PVDF membranes. Membranes were blocked for 1 hour with 5% BSA, and incubated overnight with primary antibodies (5% BSA) in 4 degrees, with Rho-GDI as the endogenous control. Membranes were washed 3Xs with TBST, incubated for 2 hours at room temperature with the appropriate secondary antibody (5% BSA or milk), and visualized using ECL.

### Quantitative PCR (qPCR)

For qPCR analysis, cells were harvested and RNA was extracted using the Maxwell RSC simplyRNA Cells kit (Promega #AS1390). cDNA synthesis was conducted with the qScript cDNA SuperMix (Quantabio #95048). qPCR was achieved using TaqMan probes to the appropriate protein of interest, with *18S* as the endogenous control.

### Chromatin Immunoprecipitation (ChIP)

1×10^6^ cells were crosslinked with 2mM DSG for 45 minutes, then with 1% formaldehyde for 25 minutes, both performed at room temperature (RT). To stop the crosslinking, glycine was added to a final concentration of 0.125M, then incubated at RT for 5 min. Cells were collected by scraping from the dishes, then washed with PBS three times. Pellets were resuspended in 0.5ml of SDS lysis buffer (1% SDS, 10mM EDTA, 50mM Tris-HCl, pH8.0)/PIC/PMSF/Sodium butyrate mix, then incubated on ice for 10 minutes. The crosslinked cellular lysates were then sonicated with a Diagnode sonicator. After sonication, samples were aliquoted into a 1.7ml tube. Tubes were centrifuged at max speed for 10 minutes at 4°C. Supernatant was then transferred to a new 1.7ml tube. To prepare chromatin immunoprecipitation sample, per 0.1ml of sonicated sample, 0.9ml of dilution buffer (50mM Tris-HCl, pH8.0, 0.167M NaCl, 1.1% Triton X-100, 0.11% sodium deoxycholate)/PIC/PMSF/Sodium butyrate mix was added, followed by antibody bound Dynabeads. The tubes were gently mixed and placed on a rocker at 4°C. The tubes were then placed in magnetic stand, inverted several times, and beads were allowed to clump. The supernatant was then discarded. Beads were flicked to resuspend and were then washed with 1X RIPA-150, 1X RIPA-500, 1X RIPA-LiCl, and 2X TE buffer (pH 8.0), for 5 minutes each on a rocker at 4°C. After each wash, the tubes were again placed on a magnetic stand and supernatant was discarded after the beads clumped. Beads were then resuspended in 200µl of Direct Elution Buffer (10mM Tris-HCl pH8.0, 0.3M NaCl, 5mM EDTA, 0.5% SDS). 1µl of RNaseA was added and incubated at 65°C to reverse crosslink. The tubes were quickly centrifuged, place on a magnetic stand, and supernatant was transferred to a new low-bind tube after beads clumped. 3µl of Proteinase K was added and incubated for 2hrs at 55°C. The sample was purified using phase lock tubes and ethanol precipitation. Samples were resuspended in 25µl of Qiagen elution buffer. DNA was amplified by real-time PCR (ABI Power SYBR Green PCR mix).

### Co-immunoprecipitation (co-IP)

Calu3 cells were incubated with either IFN-α and IFN-β, or IFN-λ for 3 hours. After IFN stimulation, cells were lysed in 1 ml of ice-cold non-denaturing lysis buffer (20 mM, Tris-HCl pH8, 137 mM NaCl, 10% glycerol, 1% Nonidet P-40, and 2 mM EDTA) supplemented with a protease inhibitor cocktail (Thermo Scientific, 78429). Whole cell extracts were incubated with either STAT1, STAT2, or control rabbit IgG antibody and protein G beads (Invitrogen, 10003D) overnight at 4 °C while rotating. Beads were washed 3 times with ice cold nondenaturing lysis buffer and boiled in 4x Laemmli Sample Buffer (Bio-Rad, #1610747) for 10 mins. For immunoblotting, samples were separated by SDS-PAGE and then transferred to PVDF membranes. Afterward, membranes were probed with antibodies against STAT1, STAT2, IRF9, and rabbit IgG.

### Immunofluorescence

Cells were plated onto round coverslips. Cells were then fixed in 4% paraformaldehyde for 15 minutes and blocked by 5% BSA for 1 hour. The cells were then incubated with primary antibodies overnight, followed by incubation with the fluorochrome-conjugated secondary anti-mouse or anti-rabbit IgG (H+L) for 1 hour at 37 °C. After staining, the slides were counterstained with DAPI (Sigma Aldrich, cat# D9542, 5 μg/ml) for 10 minutes and cover slipped with Mowiol.

### Pseudovirus SARS-Cov-2 infection

Lentivirus pseudotyped with SARS-CoV-2 Spike Protein was supplied by VectorBuilder (Catalog #: LVM-SARS-CoV-2_S(VB160426-1050cgb)-C) with Luciferase as a reporter. Following pre-exposures without and with either drugs or cytokines, AEpiC and Calu-3 cells were infected with psuedotyped SARS-CoV-2 at a multiplicity of infection of 0.1 in triplicate. Transduction efficiency of the virus was quantified 48 hours post transduction by measuring the activity of firefly luciferase in cell lysates. Luciferase signal was measured according to Dual-Glo® Luciferase Assay System - Promega Corporation.

### Gene transfer and RNA interference

shRNA expression in mammalian cells was achieved by means of lentiviral vectors. Lipofectamine 3000 was used to co-transfect transfer plasmids and packaging vectors in 293T cells. Calu-3 cells were transduced by incubation with lentiviral vector suspensions, in the presence of 8 µg/ml polybrene, for 8–12 hours. In other experiments, cDNA and small interfering RNA-expressing constructs were transiently transfected with Lipofectamine 2000 (Life Technologies) according to manufacturer’s instructions.

### Statistics

Statistical analysis used R and GraphPad Prism 8 software. At least three biologically independent samples were used to determine significance. Results are reported as mean ± SD. Non-parametric two-sided Wilcoxon rank sum tests were used to identify differentially expressed genes in all the comparisons discussed in scRNA-seq analysis. Comparisons between two groups were performed using an unpaired two-sided Student’s t test (p < 0.05 was considered significant). Comparison of multiple conditions was done with One-way or two-way ANOVA test. The Fisher’s exact test was used to compare the ratio of double positive cells between groups. Only p values of 0.05 or lower were considered statistically significant (p > 0.05 [ns, not significant], p ≤ 0.05 [∗], p ≤ 0.01 [∗∗], p ≤ 0.001 [∗∗∗], p ≤ 0.0001 [∗∗∗∗]).

## Author Contributions

F.G.G. conceived and led the study. F.G.G., Y.W. conceived the hypotheses and designed and analyzed the experiments. Y.W., S.G., H.C., S.L., E.C., Y.J., H.H.L., and F.G.G. wrote the manuscript. Y.W., S.G., H.C., S.L., E.C., Y.J., H.H.L. performed and analyzed experiments.

## Acknowledgements

This work was supported by NIH grants R35 CA197566 (Outstanding Investigator Award to F.G.G.), and by CPRIT Recruitment of Established Investigators Award RR160031 (to F.G.G.). We thank members of the Giancotti laboratory for discussions.

## Data and materials availability

All data needed to evaluate the conclusions in the paper are present in the paper or the Supplementary Materials.

---

**Fig. S1.**
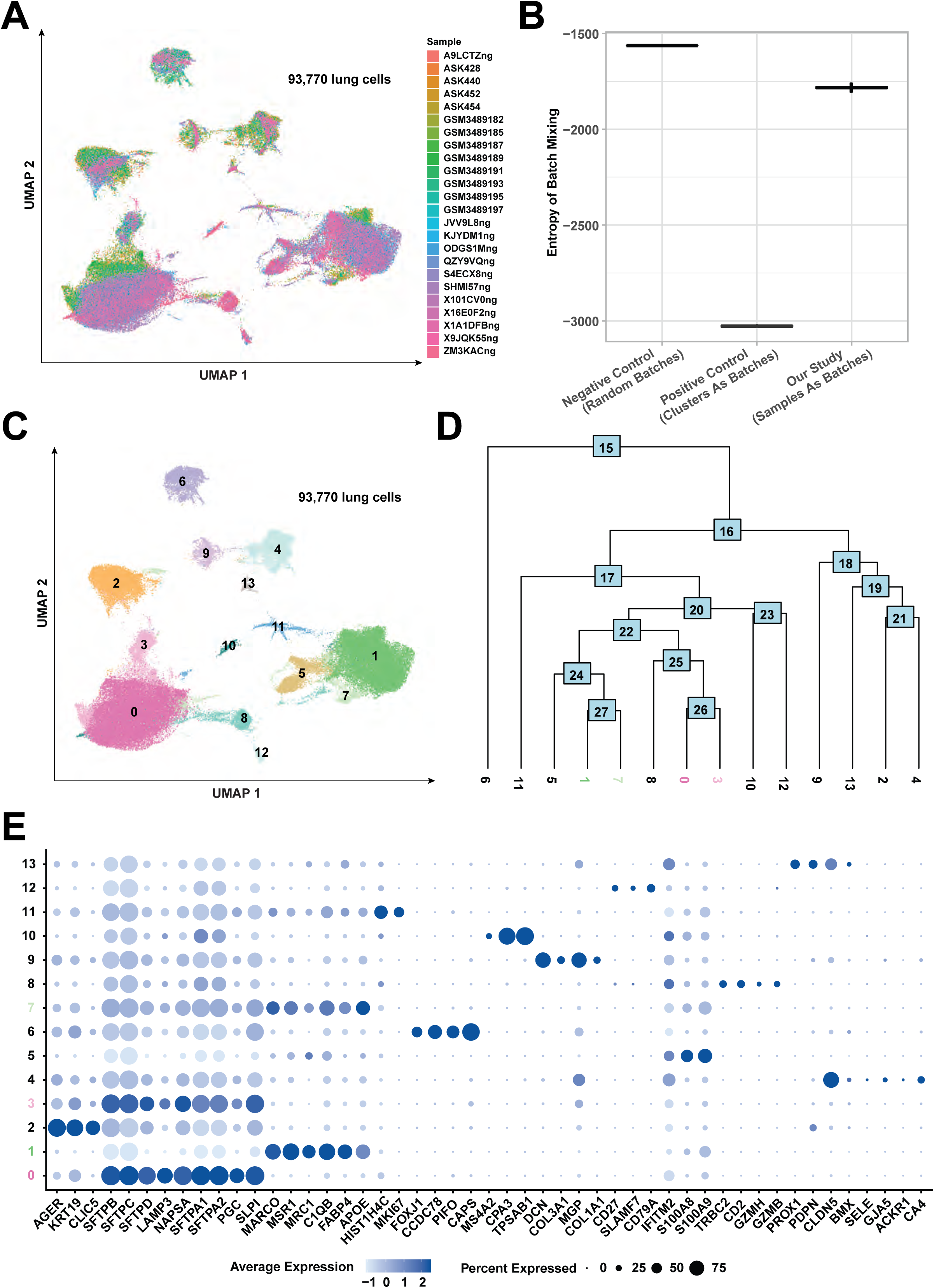
Gene expression of ACE2 and TMPRSS2 occurs largely in AT1 and AT2 epithelial cells. (A) UMAP visualization of 93,770 human lung cells colored according to sample. (B) Entropy of batch mixing for sample batches (n = 24; right); positive controls, in which clusters were assigned as batches (center); and negative controls, in which cells were assigned random batch labels in accordance with batch size distribution (left). Bars show the 25th, 50th and 75^th^ percentiles. (C) UMAP visualization of 93,770 human lung cells colored by initial cluster identity. (D) Cluster tree showing the relation between different clusters of cells. Clusters identified to be similar to each other are highlighted in green (clusters 1 and 7) or pink (clusters 0 and 3). (E) Expression of selected marker genes by cluster. Clusters identified to be similar to each other by similar expression pattern of markers are highlighted in green (clusters 1 and 7) or pink (clusters 0 and 3).

**Fig. S2.**
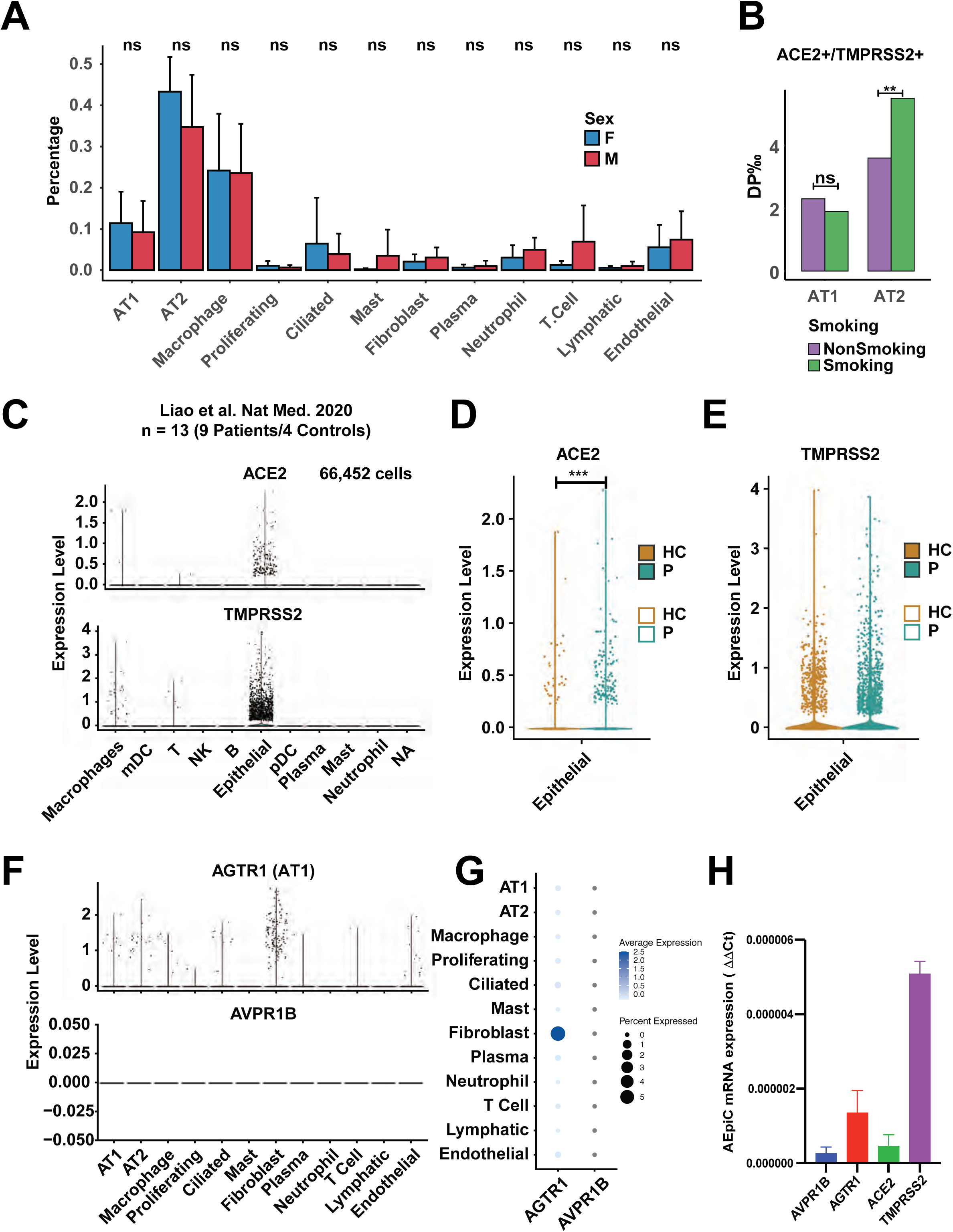
The expression level of ACE2 and TMPRSS2 is sex-related. (A) The percentages of cells of each cell type analyzed at individual sample level in males and females. (B) The ratio of ACE2+/TMPRSS2+ double positive cells in smoking and non-smoking individual groups in AT1 and AT2 cells. ** p < 0.01. (C) Normalized expression of ACE2 and TMPRSS2 in each cluster of lung cells from Liao et al. Nat Med. 2020, which includes a total of 66,452 cells from 9 patients and 4 controls. (D and E) Normalized expression of ACE2 (D) and TMPRSS2 (E) in epithelial cells in patients with COVID-19 and heathy controls. (F) Normalized expression of AGTR1 (AT1) and AVPR1B in each cluster of normal lung cells. (G) Expression of AGTR1 (AT1) and AVPR1B across all identified cell types in normal lungs. The size of the dot correlates to the percentage of cells within a cell type in which AGTR1/ AVPR1B was detected. The color encodes the average expression level. (H) Analysis of the AVPR1B, AGTR1, ACE2 and TMPRSS2 in the AEpiC cells. 5×105 AEpiC cells were harvested and subjected to RT-qPCR analysis for the indicated genes, with 18S as an endogenous control. Results are reported as mean ± SD. Comparisons between two groups were performed using an unpaired two-sided Student’s t test (p < 0.05 was considered significant). Comparison of multiple conditions was done with One-way or two-way ANOVA test. All experiments were reproduced at least three times, unless otherwise indicated.

**Fig. S3.**
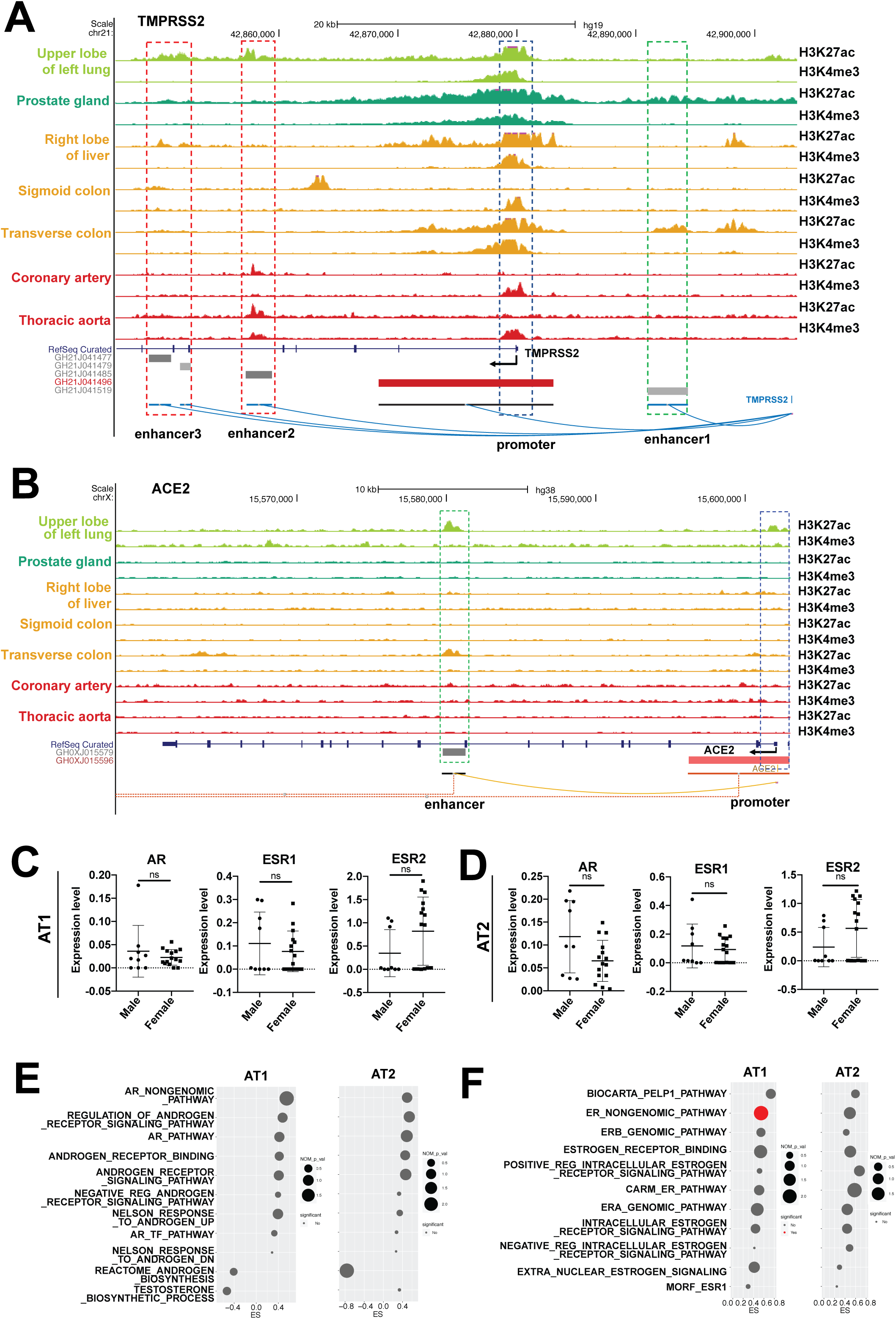
Lung-specific enhancers of TMPRSS2 are enriched in binding sites targeted by interferon transcription factors. (A) Genome browser tracks of H3K27ac and H3K4me3 ChIP-Seq data in different tissues of the expanded genomic region containing the TMPRSS2 gene. H3K27ac and H3K4me3 binding profiles are from publicly available ENCODE datasets. (B) Genome browser tracks of H3K27ac and H3K4me3 ChIP-Seq data in different tissues of the expanded genomic region containing the ACE2 gene. H3K27ac and H3K4me3 binding profiles are from publicly available ENCODE datasets. (C) Expression of AR, ESR1, and ESR2 in male and female AT1 cells. (D) Expression of AR, ESR1, and ESR2 in male and female AT2 cells. (E) GSEA gene sets involved in AR signaling enriched for genes differentially expressed in AT1 female group versus AT1 male group (left) or in AT2 female group versus AT2 male group (right). X-axis title “ES” represents the GSEA enrichment score. Y-axis represents the name of the signatures. Dot size represents the -log10 (nominal_p_value + 0.001). Dot color represents the significance. (F) GSEA gene sets involved in ER signaling enriched for genes differentially expressed in AT1 female group versus AT1 male group (left) or the AT2 female group versus AT2 male group (right). X-axis title “ES” represents the GSEA enrichment score. Y-axis represents the name of the signatures. Dot size represents the -log10 (nominal_p_value + 0.001). Dot color represents the significance.

**Fig. S4.**
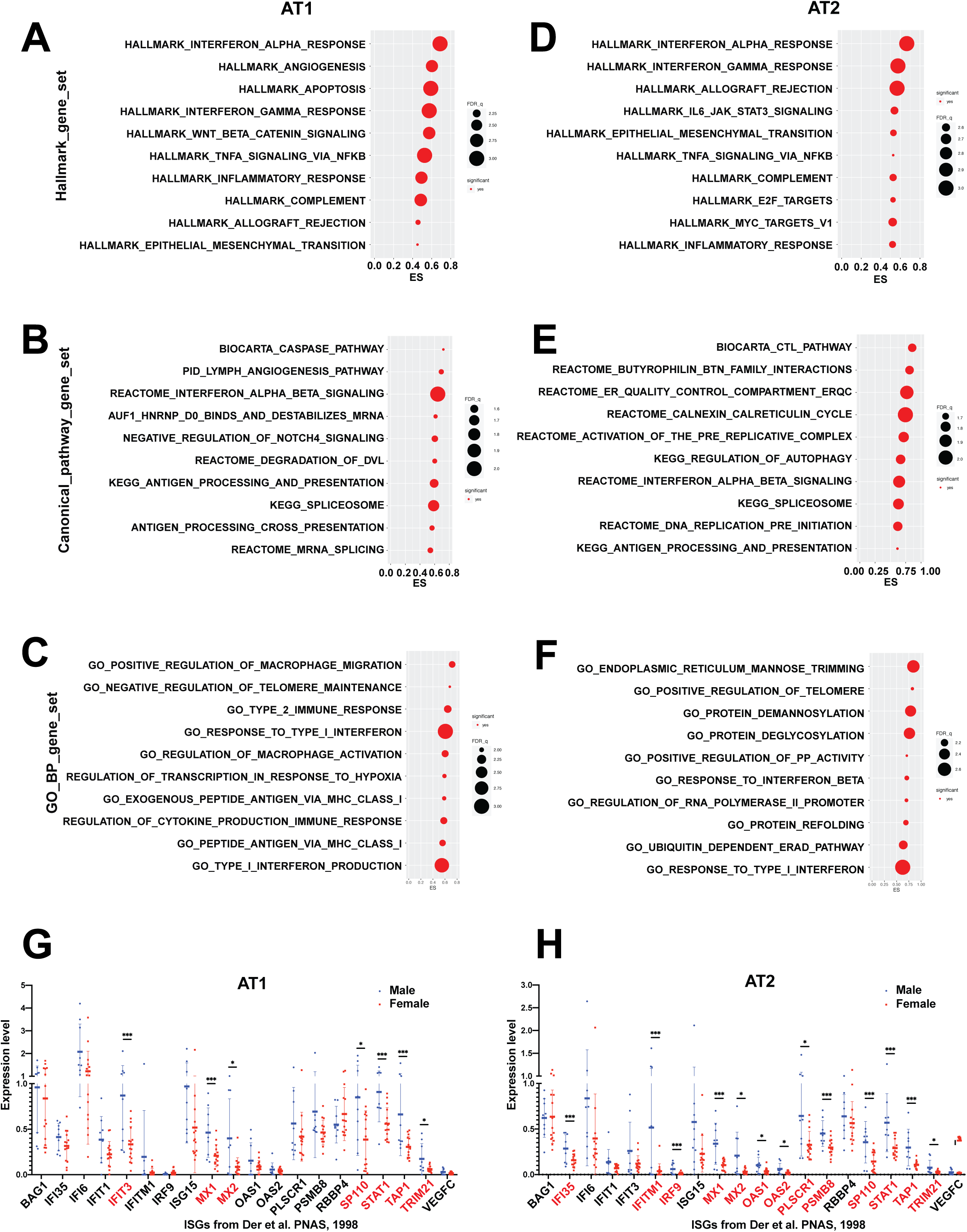
The activity of interferon signaling pathways is higher in male lung AT1 and AT2 cells than in female lung AT1 and AT2 cells. (A) GSEA signatures from hallmark gene sets enriched for genes differentially expressed in AT1 female group versus AT1 male group. X-axis title “ES” represents the GSEA enrichment score. Y-axis represents the name of the signatures. Dot size represents the -log10 (FDR_q_value + 0.001). Dot color represents the significance. (B) GSEA signatures from canonical pathway gene sets enriched for genes differentially expressed in AT1 female group versus AT1 male group. X-axis title “ES” represents the GSEA enrichment score. Y-axis represents the name of the signatures. Dot size represents the -log10 (FDR_q_value + 0.001). Dot color represents the significance. (C) GSEA signatures from GO biological process gene sets enriched for genes differentially expressed in AT1 female group versus AT1 male group. X-axis title “ES” represents the GSEA enrichment score. Y-axis represents the name of the signatures. Dot size represents the -log10 (FDR_q_value + 0.001). Dot color represents the significance. (D) GSEA signatures from hallmark gene sets enriched for genes differentially expressed in AT2 female group versus AT2 male group. X-axis title “ES” represents the GSEA enrichment score. Y-axis represents the name of the signatures. Dot size represents the -log10 (FDR_q_value + 0.001). Dot color represents the significance. (E) GSEA signatures from canonical pathway gene sets enriched for genes differentially expressed in AT2 female group versus AT2 male group. X-axis title “ES” represents the GSEA enrichment score. Y-axis represents the name of the signatures. Dot size represents the -log10 (FDR_q_value + 0.001). Dot color represents the significance. (F) GSEA signatures from GO biological process gene sets enriched for genes differentially expressed in AT2 female group versus AT2 male group. X-axis title “ES” represents the GSEA enrichment score. Y-axis represents the name of the signatures. Dot size represents the -log10 (FDR_q_value + 0.001). Dot color represents the significance. (G). Expression of interferon-stimulated genes in male (blue) and female (red) AT1 cells. Genes showing significant difference between male and female group are highlighted in red. (H) Expression of interferon-stimulated genes in male (blue) and female (red) AT2 cells. Genes showing significant difference between male and female group are highlighted in red.

**Fig. S5.**
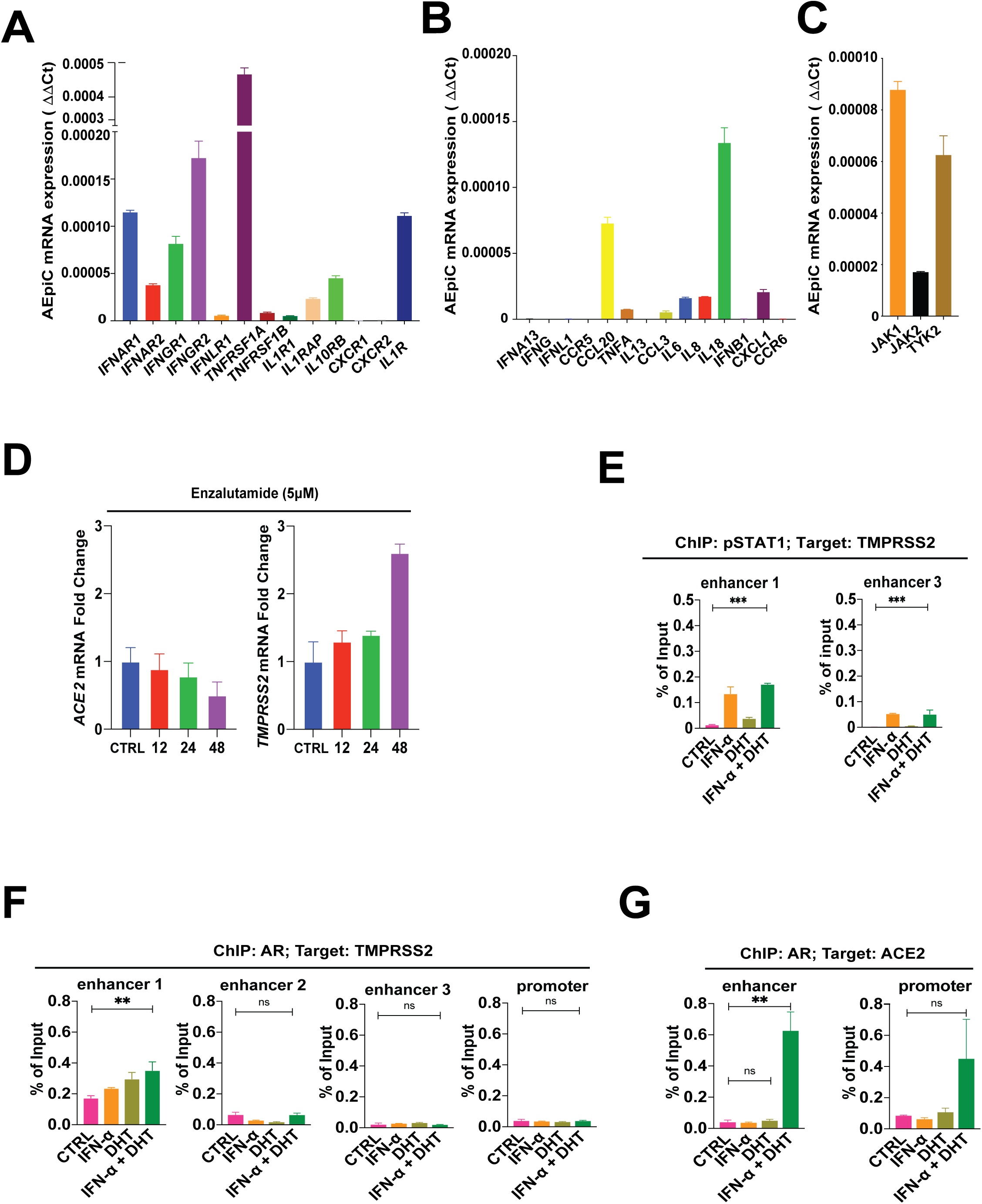
Interferon-stimulated JAK/STAT signaling, instead of AR-dependent signaling, transactivates the expression of SARS-CoV2 receptors, TMPRSS2 and ACE2, in human pulmonary alveolar epithelium. (A, B, C) Analysis of the signaling components of the IFN-JAK-STAT pathway and COVID-19 advanced lung disease-related cytokines and its receptors in the AEpiC cells. 5×105 AEpiC cells were harvested and subjected to RT-qPCR analysis for the indicated genes, with 18S as an endogenous control. (D) The change of SARS-CoV-2 receptors mRNA expression level during Enzalutamide treatment. Enzalutamide treatment (5 μM) of AEpiC cells for the indicated times. RT-qPCR was performed for ACE2 (left) and TMPRSS2 (right), with 18S as an endogenous control. (E) Enrichment of p-STAT1 on enhancer 1 and 3 of TMPRSS2 from ChIP-qPCR in control. 1×106 AEpiC cells were either treated with DHT alone (10 nM, 24 hours), IFN⍺ alone (20 nM, 16 hours), or in combination. Cells were lysed in a non-denaturing lysis buffer and subjected to ChIP of p-STAT1. (F and G) Enrichment of AR on the enhancer and the promoter regions of TMPRSS2 and ACE2 from ChIP-qPCR in control. 1×106 AEpiC cells were either treated with DHT alone (10 nM, 24 hours), IFN⍺ alone (20 nM, 16 hours), or in combination. Cells were lysed in a non-denaturing lysis buffer and subjected to ChIP of AR. Results are reported as mean ± SD. Comparisons between two groups were performed using an unpaired two-sided Student’s t test (p < 0.05 was considered significant). Comparison of multiple conditions was done with One-way or two-way ANOVA test. All experiments were reproduced at least three times, unless otherwise indicated.

**Fig. S6.**
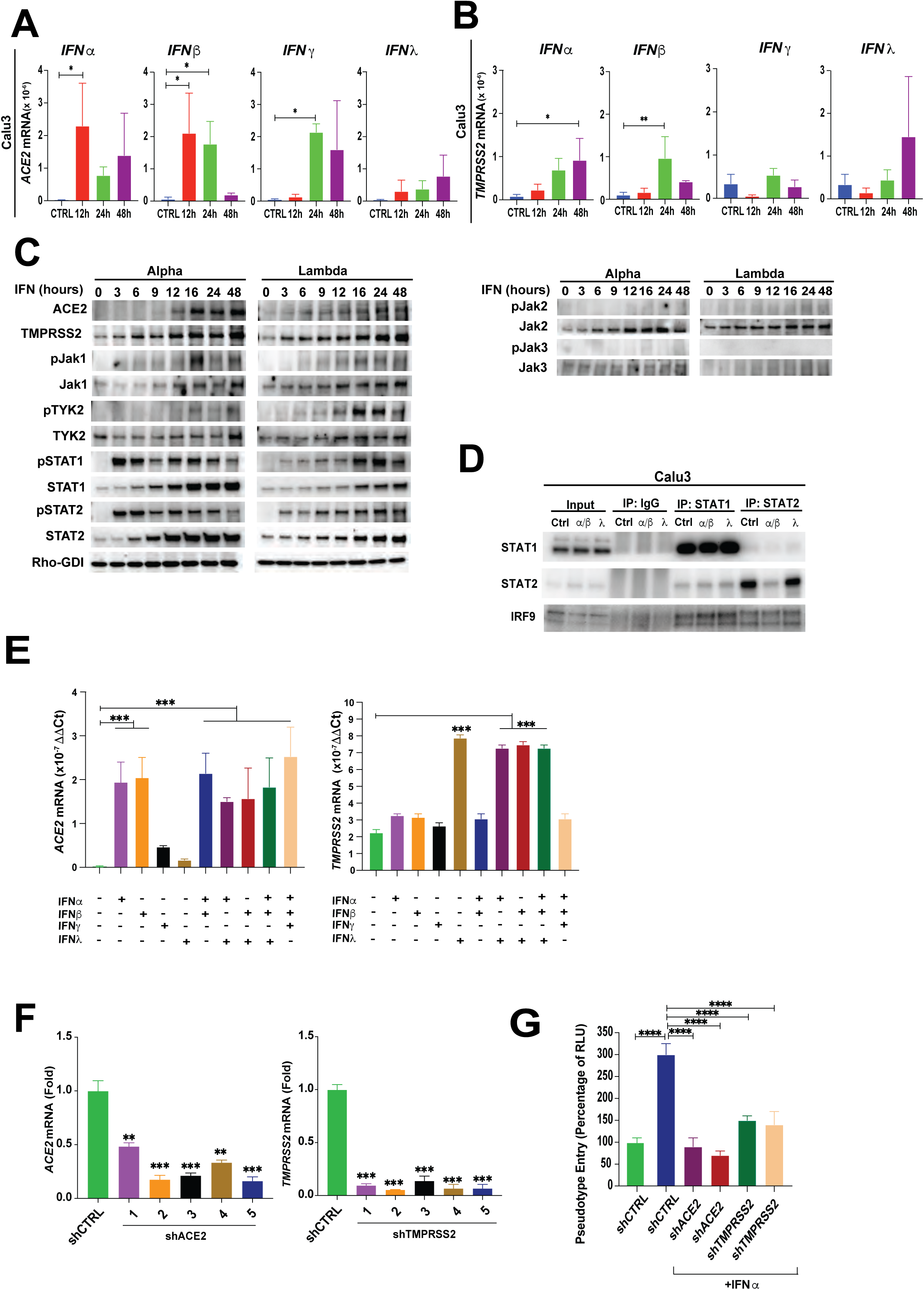
Interferons a/b and l conspire to upregulate the expression of ACE2 and TMPRSS2 and induce robust viral entry. (A) Calu-3 were treated with either PBS, 10 nM IFN⍺, 10 nM IFNβ, 10ng/ml IFNɣ, or 1 μg/ml IFNλ. ACE2 mRNA expression measured with RT-qPCR, with 18S as an endogenous control. (B) Calu-3 were treated with either PBS, 10 nM IFN⍺, 10 nM IFNβ, 10ng/ml IFNɣ, or 1 μg/ml IFNλ. TMPRSS2 mRNA expression measured with RT-qPCR, with 18S as an endogenous control. (C) Activation of JAK-STAT signaling pathway and the expression of SARS-CoV-2 receptors in pulmonary alveolar epitheliums during Type I and Type III interferon stimulation. AEpiC cells were treated with 20 nM IFN⍺ (left) or 1 μg/ml IFNλ (right) for the indicated times. Total lysates were subjected to western blot with the indicated antibodies, with GDI as an internal loading control. Source data-Supplementary Figure 6C-1. Source data-Supplementary Figure 6C-2. (D) Calu-3 control cells or cells treated with 10 nM IFN⍺ and 10nM IFNβ, or 1 μg/ml IFNλ for 3 hours were subjected to Co-IP. Either input (left), IgG, STAT1, or STAT2 (right) was immunoprecipitated, and total lysates were subjected to western blot with the indicated antibodies. Source data-Supplementary Figure 6D. (E) SARS-CoV-2 receptors mRNA expression level in response to different combinations of interferon stimulation. AEpiC cells were treated with 10 nM IFN⍺, 10 nM IFNβ, 10 ng/ml IFNɣ or 1 ug/ml IFNλ, alone or in different combinations as indicated in the graph. RT-qPCR was performed for ACE2 (Left) and TMPRSS2 (Right), with 18S as an endogenous control. (F) Calu-3 cells subjected to shRNA of ACE2 (left) or TMPRSS2 (right). 5 shRNAs used per gene. ACE2 and TMPRSS2 mRNA expression measured with RT-qPCR, with 18S as an endogenous control. (G) Calu-3 cells silenced for control, ACE2 or TMPRSS2, and treated either with PBS or 10 nM IFN⍺ for 16 hours before viral infections. The transduction efficiency of the virus was quantified 48 hours post-transduction by measuring the activity of firefly luciferase in cell lysates. Results are reported as mean ± SD. Comparisons between two groups were performed using an unpaired two-sided Student’s t test (p < 0.05 was considered significant). Comparison of multiple conditions was done with One-way or two-way ANOVA test. All experiments were reproduced at least three times, unless otherwise indicated.

**Fig. S7.**
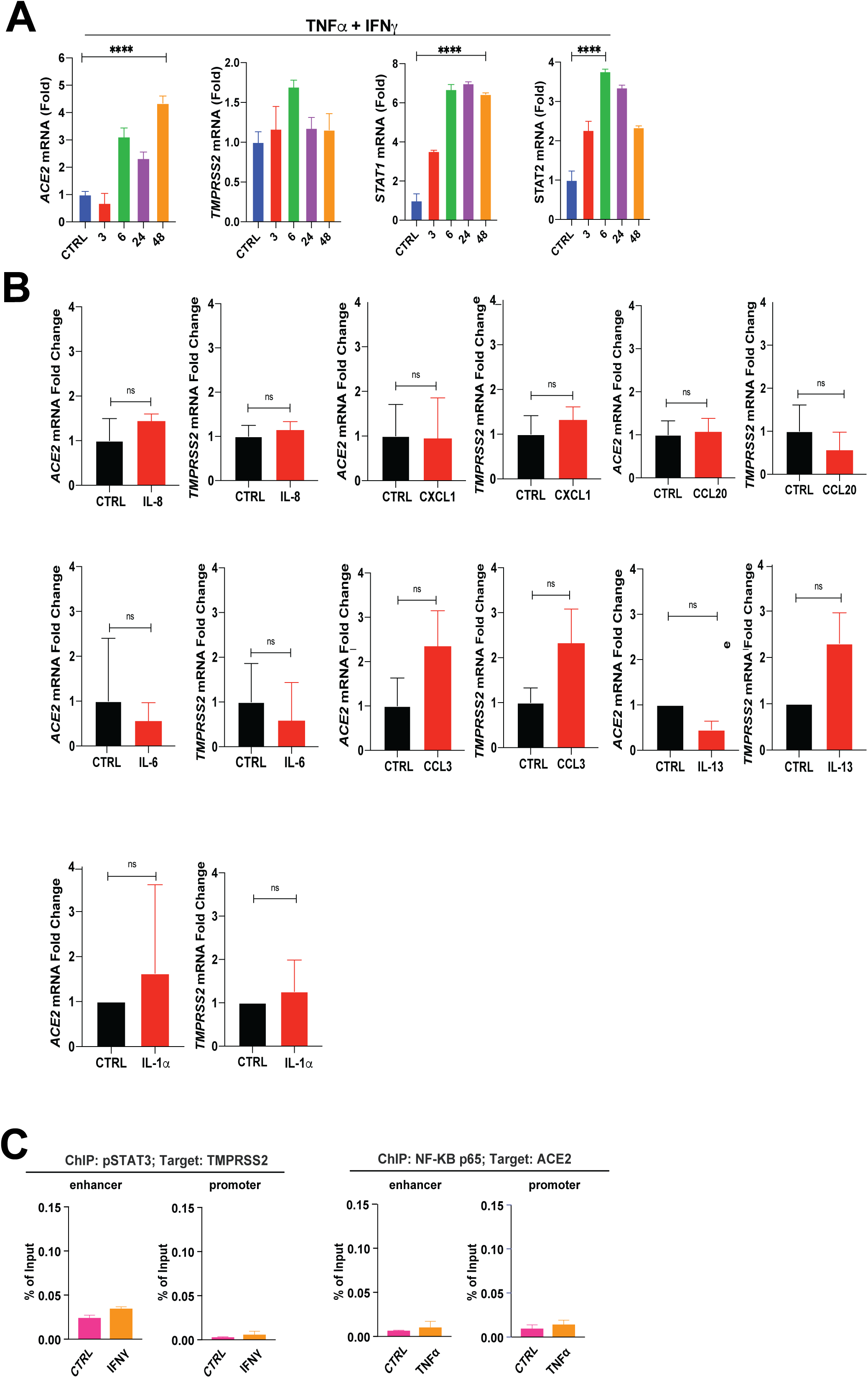
TNF⍺ models severe SARS-COV2 disease and cytokine storm in pulmonary alveolar epithelial cells. (A) AEpiC cells treated with PBS, 10 ng/ml TNF⍺ plus 10 ng/ml IFNɣ for 24 hours. ACE2, TMPRSS2, STAT1, and STAT2 mRNA expression measured with RT-qPCR, with 18S as an endogenous control. (B) AEpiC cells were treated with different cytokines (all 10 ng/ml except 5ng/ml for IL-6) for 24 hours. ACE2 and TMPRSS2 mRNA expression measured with RT-qPCR, with 18S as an endogenous control. (C) Enrichment of p-STAT3 on the enhancer and the promoter of ACE2 from ChIP-qPCR in control and IFNɣ of AEpiC cells. Enrichment of NF-kB p65 on the enhancer and the promoter of TMPRSS2 gene from ChIP-qPCR in control and TNF⍺ of AEpiC cells. Results are reported as mean ± SD. Comparisons between two groups were performed using an unpaired two-sided Student’s t test (p < 0.05 was considered significant). Comparison of multiple conditions was done with One-way or two-way ANOVA test. All experiments were reproduced at least three times, unless otherwise indicated.

## Supplemental material

**Supplementary Table 1.** Resources table

| **<u>Reagent/Resource</u>** | **<u>Source</u>** | **<u>Identifier</u>** |
| --- | --- | --- |

| **Antibodies** |  |  |
| --- | --- | --- |
| ACE2 | R&D Systems | AF933 |
| TMPRSS2 | Abcam | ab92323 |
| Jak2 | Cell Signaling | 3230 |
| p-Jak2 | Cell Signaling | 3776 |
| STAT1 | Cell Signaling | 14994 |
| p-STAT1 | Cell Signaling | 9167 |
| STAT2 | Cell Signaling | 72604 |
| p-STAT2 | Cell Signaling | 88410 |
| STAT3 | Cell Signaling | 9139 |
| p-STAT3 | Cell Signaling | 9145 |
| Jak1 | Cell Signaling | 3332 |
| p-Jak1 | Cell Signaling | 3331 |
| TYK2 | Cell Signaling | 9312 |
| p-TYK2 | Cell Signaling | 9321 |
| IRF9 | Santa Cruz | Sc-514648 |
| Jak3 | Cell Signaling | 3775 |
| p-Jak3 | Cell Signaling | 5031 |
| Rho-GDI | Santa Cruz | Sc-373724 |
| Rabbit IgG | Cell Signaling | 3900 (IP), 7074 (WB) |
| **Chemicals, Cell Lines, and Recombinant Proteins** |  |  |
| DMEM | ThermoFisher Scientific | 11965-092 |
| Alveolar Epithelial Cell Medium | FisherScientific | NC9028239 |
| Penicillin G-Streptomycin | Corning | 30004CI |
| Trypsin-EDTA (0.25%) | ThermoFisher Scientific | 2530054 |
| FBS | ThermoFisher Scientific | A4766801 |
| Calu-3 | ATCC | HTB-55 |
| LNCap | ATCC | CRL-1740 |
| AEpiC | FisherScientific | 501049330 |
| rhIL-8 | R&D Systems | 208-IL |
| rhIL-6 | R&D Systems | 206-IL |
| rhCCL20/MIP-3a | R&D Systems | 360-MP |
| rhCXCL1/GROa | R&D Systems | 275-GR |
| rhCCL3/MIP-1a | R&D Systems | 270-LD |
| rhIL-1a | Fisher | 200LA |
| rhIL-13 | Fisher | 213ILB |
| rhTNF-a | R&D Systems | 210-TA |
| rhIFN-lambda | R&D Systems | 1598-IL |
| rhIFN-alpha | Sigma-Aldrich | IF007 |
| rhIFN-beta | Sigma-Aldrich | IF014 |
| rhIFN-gamma | R&D Systems | 285-IF |
| **TaqMan Probes** |  |  |
| ACE2 | Fisher | Hs01085333_m1 |
| TMPRSS2 | Fisher | Hs01122322_m1 |
| STAT1 | Fisher | Hs01013996_m1 |
| STAT2 | Fisher | Hs01013115_g1 |
| 18S | Fisher | Hs99999901_s1 |
| **Reagents** |  |  |
| dsDNA naked | Invitrogen | tlrl-patn |
| Poly(I:C) LMW | Invitrogen | tlrl-picw |
| LPS | Invitrogen | tlrl-pb5lps |

| Software and Algorithms |  |  |
| --- | --- | --- |
| R | R Development Core Team, 2016 | https://www.r-project.org/ |
| Seurat v3.2.0 | Stuart et al., 2019 | https://satijalab.org/seurat/ |
| Harmony | Korsunsky et al., 2019 | https://github.com/immunogenomics/harmony |
| ENCODE | ENCODE Consortium | https://www.encodeproject.org |
| UCSC Genome Browser | Kent et al., 2002 | https://genome.ucsc.edu |
| GeneHancer<br>Regulatory Elements and Gene Interactions | Fishilevich et al., 2017 | https://genome.ucsc.edu/cgi-bin/hgTrackUi?db=hg19&g=geneHancer |
| GSEA 20.0.5 | Tamayo, et al., 2005 | https://www.gsea-msigdb.org/gsea/index.jsp |

**Supplementary Table 3.** The Encyclopedia of DNA Elements (ENCODE) data sets

| Experiment | File download | Target | Tissue |
| --- | --- | --- | --- |
| ENCSR153NDQ | ENCFF250INB | H3K4me3 | prostate gland male adult (37 years) |
| ENCSR841AJO | ENCFF121EZK | H3K27ac | prostate gland male adult (37 years) |
| ENCSR074WIB | ENCFF602GRK | H3K4me3 | upper lobe of left lung male adult (37 years) |
| ENCSR505YFA | ENCFF027NNY | H3K27ac | upper lobe of left lung male adult (37 years) |
| ENCSR344TLI | ENCFF249QWR | H3K4me3 | right lobe of liver female adult (53 years) |
| ENCSR981UJA | ENCFF599LSB | H3K27ac | right lobe of liver female adult (53 years) |
| ENCSR807XUB | ENCFF898XDY | H3K27ac | sigmoid colon male adult (37 years) |
| ENCSR960AAL | ENCFF378PGI | H3K4me3 | sigmoid colon male adult (37 years) |
| ENCSR640XRV | ENCFF284LUF | H3K27ac | transverse colon male adult (37 years) |
| ENCSR813ZEY | ENCFF604IEC | H3K4me3 | transverse colon male adult (37 years) |
| ENCSR343ZOV | ENCFF297RMA | H3K4me3 | coronary artery female adult (53 years) |
| ENCSR443UYU | ENCFF172MCO | H3K27ac | coronary artery female adult (53 years) |
| ENCSR015GFK | ENCFF772QFT | H3K27ac | thoracic aorta male adult (37 years) |
| ENCSR930HLX | ENCFF712GED | H3K4me3 | thoracic aorta male adult (37 years) |

